# The Stmn1-lineage contributes to acinar regeneration but not to neoplasia upon oncogenic Kras expression

**DOI:** 10.1101/2025.03.18.643944

**Authors:** Shakti Dahiya, Jorge R. Arbujas, Arian Hajihassani, Sara Amini, Michael Wageley, Klara Gurbuz, Zhibo Ma, Celina Copeland, Mohamed Saleh, George K. Gittes, Bon-Kyoung Koo, Kathleen E. DelGiorno, Farzad Esni

**Affiliations:** Department of Surgery, Division of Pediatric General and Thoracic Surgery, Children’s Hospital of Pittsburgh, University of Pittsburgh Medical Center, Pittsburgh, PA 15244, United States; Department of Cell and Developmental Biology, Vanderbilt University, Nashville, TN 37232, United States; Department of Pediatric, Children’s Hospital of Pittsburgh, University of Pittsburgh Medical Center, Pittsburgh, PA 15244, United States; Institute of Molecular Cell Biotechnology of Austrian Academy of Science (IMBA), 1030 Vienna, Austria; Vanderbilt Digestive Disease Research Center, Vanderbilt University Medical Center, Nashville, TN; Vanderbilt Ingram Cancer Center, Vanderbilt University Medical Center, Nashville, TN; Center for Computational Systems Biology, Vanderbilt University Medical Center, Nashville, TN; Department of Developmental Biology, University of Pittsburgh, Pittsburgh, PA 15244, United States; UPMC Hillman Cancer Center, Pittsburgh, PA, 15123, United States; McGowan Institute for Regenerative Medicine, University of Pittsburgh, Pittsburgh, PA 15244, United States

**Keywords:** Pancreas, Stmn1, Acinar cells, Regeneration, ADM, Neoplasia

## Abstract

**BACKGROUND & AIMS:** The exocrine pancreas has a limited regenerative capacity, but to what extent all acinar cells are involved in this process is unclear. Nevertheless, the heterogenous nature of acinar cells suggests that cells exhibiting higher plasticity might play a more prominent role in acinar regeneration. In that regard, *Stmn1*-expressing acinar cells have been identified as potential facultative progenitor-like cells in the adult pancreas. Here, we studied Stmn1-progeny under physiological conditions, during regeneration, and in the context of *Kras^G12D^* expression.

**METHODS:** We followed the fate of Stmn1-progenies both under baseline conditions, following caerulein-induced acute or chronic pancreatitis, pancreatic duct ligation, and in the context of *Kras^G12D^* expression.

**RESULTS:** The Stmn1-lineage contributes to baseline acinar cell turnover under physiological conditions. Furthermore, these cells rapidly proliferate and repopulate the acinar compartment in response to acute injury in an ADM-independent manner. Moreover, acinar regeneration during chronic pancreatitis progression is in conjunction with a decline in the proliferative capacity of the Stmn1-lineage. Interestingly, newly generated acinar cells display increased susceptibility to additional injury during recurrent acute pancreatitis (RAP). Finally, given their inability to form ADMs, the Stmn1-lineage fails to form PanINs upon oncogenic *Kras* expression.

**CONCLUSIONS:** Our findings establish the Stmn1-lineage as a pivotal subpopulation for acinar regeneration. The ability of these cells to restore acinar tissue in an ADM-independent manner distinguishes them as a critical regenerative population. This study presents a new paradigm for acinar regeneration and repair in the context of pancreatitis and neoplasia.

## Introduction

Acute Pancreatitis (AP) develops as the result of pancreatic exocrine injury, premature activation of digestive enzymes and subsequent cellular death and release of pro-inflammatory cytokines^1, 2^. Chronic Pancreatitis (CP) is a condition characterized by progressive pancreatic inflammation, fibrosis, acinar atrophy, and severe pain^3, 4^. In progressive, predictive models of chronic pancreatitis (CP), multiple episodes of AP, or Recurrent Acute Pancreatitis (RAP) may lead to CP^3^. In experimental models of AP, 25-50% of the pancreatic parenchyma is destroyed. However, the obliterated portions are almost completely regenerated within a week after injury. One of the earliest events following exocrine injury is acinar-to-ductal metaplasia (ADM), a reversible process induced by cytokines and chemokines released either by injured acini or infiltrating innate immune cells, mainly macrophages^3, 5–10^. It is generally accepted that acinar regeneration is through ADMs, which entails dedifferentiation of acinar cells into duct-like cells, followed by proliferation of metaplastic ducts^11–13^. Tissue recovery in pancreatitis is orchestrated by a delicate balance between inflammation and acinar regeneration. Thus, concomitant with the resolution of inflammation in AP, these metaplastic ducts are thought to re-differentiate into acinar cells^13, 14^. Accordingly, the persistence of those pro-inflammatory signals in CP may prevent re-differentiation of ADMs into acinar cells in CP, making it an irreversible disease with no apparent acinar regeneration. Nevertheless, to what extent the acinar compartment in CP could regain its regenerative capacity upon elimination of inflammation and fibrosis is unknown.

CP is also considered as a risk factor for pancreatic ductal adenocarcinoma (PDAC). PDAC arises from noninvasive precursor lesions, among which pancreatic intraepithelial neoplasia (PanIN) is perhaps the most common and extensively examined. Studies using mouse genetic models show that upon oncogenic KRAS-induced ADM, these metaplastic structures become irreversible and act as precursors to neoplastic lesions^15, 16^. Given the involvement of ADM in acinar regeneration and its role as precursor to PanINs, pancreatic neoplasia has been popularly defined as a regenerative process highjacked by oncogenic KRAS.

The degree of heterogeneity that exists within acinar cells in the adult mouse and human pancreas is well accepted^17–19^. Nevertheless, the potential contribution of different subpopulations to acinar tissue homeostasis and regeneration is uncertain, as previous studies present two opposing models for acinar regeneration. One model proposes that despite existing heterogeneity, all acinar cells in the adult pancreas can equally contribute to tissue maintenance^20^. Another model argues for the presence of a distinct subset of acinar cells with progenitor signatures that plays a crucial role in both homeostasis and regeneration^18, 21–23^. Acinar cells expressing *Bmi1*, *Dclk1*, *Tert* or *Tff2* are all involved in tissue homeostasis^18, 21–23^. However, these subpopulations behave differently during regeneration and carcinogenesis. The Tff2-lineage (defined as transient amplifying^24^) regresses during injury and is refractory to oncogenic Kras, whereas the other lineages (defined as facultative^24^) expand following injury and can form neoplastic lesions upon exposure to *Kras^G12D^*.

In that regard, Stathmin-1 (Stmn1)-expressing acinar cells have been suggested as potential facultative progenitor-like cells in the adult pancreas^17^. STMN1 is a cytoplasmic protein that mediates formation of the mitotic spindle during cell division by regulating microtubule polymerization^25–28^. STMN1 has also been implicated in the regulation of cell proliferation through effects on cell-cycle factors^29^. Accordingly, singe cell RNA-seq analysis by us and others^17^ uncovered a subpopulation of STMN1^+^ acinar cells with retained proliferative activity in both mouse and human pancreas. Here, we aimed to study the contribution of the Stmn1-lineage in acinar homeostasis and during regeneration and neoplasia *in vivo*.

## Methods

### Study Approval

Mice used in these studies were maintained according to protocols approved by the University of Pittsburgh, the Salk Institute for Biological Studies, or Vanderbilt University Institutional Animal Care and Use Committees.

### Mice

The Rosa26*^CAGTomato^* (Gt(Rosa)26Sor*^tm^*^14^*^(CAG-td-Tomato)Hze^*^)30^, Ptf1a*^CreERTM;^* Rosa*^EYFP^*^/+ 31^, and wild type C57bl/6 mice were purchased from The Jackson Laboratory. The *Stmn1-P2A-eGFP-IRES-CreERT2* strain was generated in Dr. Bon-Kyoung Koo laboratory^32^. LSL-Kras*^G12D/+^* mice have been previously described in detail^33^. Both male and female mice were used for the experiments, unless otherwise stated. All mice used in this study were between 8 and 12 weeks of age at the beginning of the experiments. Mice were housed 4-5/cage and maintained in an environment at 20-23 °C and 30-70% humidity with ad libitum access to regular chow (Prolabs IsoPro® RMH 3000, LabDiet, 30005737-220) and water under a 12-hour light/12-hour dark cycle with the lights on from 7am until 7pm.

### Genotyping

Genotyping was performed either with Transnetyx or using KAPA Express Extract Kit (Fischer Scientific, 50-196-5275) and GoTaq® Master Mix (Promega, M7123). Primers per mouse line were used as follow: **Stmn1CreERT2:** Forward: 5’- ACCTGAAGATGTTCGCGATTATCT -3’, Reverse: 5’- ACCGTCAGTACGTGAGATATCTT -3’; **R26-TdTomato:** Wild Type Forward:5’-AAG GGA GCT GCA GTG GAG TA-3’, Wild Type Reverse:5’-CCG AAA ATC TGT GGG AAG TA-3’, tdTomato Mutant Forward:5’- CTG TTC CTG TAC GGC ATG G-3’, tdTomato Mutant Reverse:5’-GGC ATT AAA GCA GCG TAT CC- 3’;

### Tamoxifen treatment

For Cre-recombinase activation, Stmn1CreERT2;R26*^Tom^*mice were gavaged once daily with 5 mg Tamoxifen (Sigma-Aldrich, T5648) dissolved in corn oil for 3 days, as described elsewhere^34^. Ptf1a*^CreERTM^*; Rosa*^EYFP/+^* mice were treated as previously described^19^.

### Caerulein induced acinar injury

Pancreatic injury in mice was induced by a standard pancreatitis induction method using the cholecystokinin (CCK) analog caerulein (Sigma-Aldrich, C9026) at a hyper-stimulatory dose of 125 μg/kg. Mice received 8 hourly intraperitoneal cearulein injections on two consecutive days (acute pancreatitis). To induce chronic pancreatitis, mice received repeated acute pancreatitis regiment for four or eight weeks. Ptf1a*^CreERTM^*; Rosa*^EYFP/+^* mice were treated with two weeks of caerulein (250 μg/kg) twice daily, for 5 days. Ptf1a*^CreERTM^*; LSL-Kras*^G12D^*;Rosa*^EYFP/+^* mice were treated with caerulein (250 μg/kg) once daily, for 5 days.

### Diphtheria toxin treatment

Stmn1CreERT2;R26*^DTR/Tom^*, or Stmn1CreERT2;R26*^DTR^* mice treated intraperitoneally with 0.5 ng/g body weight of DT for 2 consecutive days, as described previously^35^.

### Pancreatic duct ligation

Pancreatic duct ligation was performed by standard procedure. Briefly, mice were anesthetized using isoflurane, and a midline laparotomy was performed to expose the pancreas. The main pancreatic duct was identified at the junction between the head and body of the pancreas. Using microsurgical forceps and fine sutures (6-0), the pancreatic duct was ligated, ensuring minimal disruption to surrounding blood vessels.

### Tissue processing and Immunostaining

Tissue processing, and immunostaining were performed as previously described^35^. The following antibodies were used: goat anti-Amylase (1:250, Santa Cruz, sc-12821); rabbit anti-Amylase (1:1000, Sigma-Aldrich, A8273); rat anti F4/80 (1:100, Abcam, ab6640); rabbit anti-Claudin-18 (1:100, Thermo Fisher Scientific, 38-8000); DBA FITC-conjugated (1:100, Vector Laboratories, FL1031-2); rat-anti E-cadherin (1:200, Thermo Fisher Scientific, 13-1900); goat anti-GFP (1:1000, Abcam, ab6673)rabbit anti-PCNA (1:1000, Abcam, ab92552); rabbit anti-STMN1 (1:2000, Abcam, ab52630); rat anti Ki67 (1:200, Thermo Fisher Scientific, 14-5698-82).

The following secondary antibodies used for immunostaining were purchased from Jackson ImmunoResearch Laboratories: biotin-conjugated anti-rabbit (1:500, 711-066-152), biotin-conjugated anti-rat (1:500, 712-066-153), biotin conjugated anti-guinea pig (1:500, 706-065-148), biotin-conjugated anti-goat (1:250, 705-065-147); Cy2-conjugated streptavidin (1:500, 016-540-084); Cy3-conjugated streptavidin (1:500, 016-160-084); Cy5-conjugated streptavidin (1:100, 016-600-084); Cy2-conjugated anti-guinea pig (1:300, 706-545-148), Cy3-conjugated anti-guinea pig (1:300, 706-166-148), Cy2- conjugated anti-rabbit (1:300, 711-485-152), Cy3-conjugated anti-rabbit (1:300, 711-165-152), Cy2- conjugated anti-rat (1:300, 712-545-153), Cy3-conjugated anti-rat (1:300, 712-166-150), Cy2- conjugated anti-goat (1:300, 705-545-147) and Cy3-conjugated anti-goat (1:300, 705-165 147). Additional secondaries: TSA Vivid fluorophore kit 650 (1:500, Tocris Bioscience, 7527), TSA Vivid fluorophore kit 570 (1:750, Tocris Bioscience, 7526), anti-goat 488 (1:500, Invitrogen, A32814). Prolong gold antifade reagent with DAPI (Invitrogen, P36931) was used for mounting.

### Analysis of published single-cell RNA sequencing dataset

Processed count matrices for scRNA-seq datasets from Ma et.al., Schlesinger et al., and Hendley et al., were downloaded from the Gene Expression Omnibus (GEO) database (accession numbers GSE172380, GSE141017, GSE159343) and analyzed as previously described^19, 36, 37^. Briefly, low quality cells were filtered on read counts, the number of genes expressed, and the ratio of mitochondrial reads following the thresholds described in the respective publications. Filtered gene count matrices were log-normalized, and the top 2000 variable features were further scaled prior to dimension reduction by PCA and being embedded in UMAP using the R package Seurat. Seurat cell clusters were labeled with major cell types using marker genes provided by authors.

### Fluorescent Imaging

Imaging of pancreatic tissue sections was performed using a Leica Dmi8 fluorescent light microscope at 10X, 20X or 63x objectives using LASX software or an Olympus VS200 slide scaner. The microscope is equipped with 405, 488, 568 and 647nm filters.

### Confocal imaging

Sections were imaged using a Leica Stellaris 5 confocal laser scanning microscope at 20X or 63X objectives using LASX software. Final figures were composed using Adobe Photoshop.

### Measurements of acinar area

Sections were collected serially so that each slide would contain semi-adjacent sections across the entire tissue. H&E sections were scanned using the EVOS m7000 Software using the 10x objectives. ImageJ was then employed to measure the total pancreatic area and the acinar area, evaluating the head, body and tail regions of various experimental groups, as described in the text.

### Quantification analysis

To calculate the percentage Tom^+^/Amy^+^, Tom^+^/STMN1^+^, Tom^+^/Ki67^+^ cells mice, whole pancreata were sectioned, and sections separated by 200-300 μm were stained for amylase, STMN1 or Ki67. Data were obtained by analyzing 10x images of pancreata cross sections. Captured images of islets were analyzed using ImageJ software.

### Statistical analysis

Comparisons between 2 groups were made using unpaired, two-tailed t-test as indicated. Statistical analysis was performed using GraphPad Prism 9 software (GraphPad software version 9.2.0, San Diego, CA). Unless specified, data in the text, table and figures is expressed as a mean ± standard deviation (SD).

## Results

### The Stmn1-lineage contributes to acinar tissue homeostasis

To study the facultative capacity of *Stmn1*-expressing cells, we first followed the fate of Stmn1-progeny under physiological conditions in Stmn1-CreERT2;R26*^Tom^* (SC*^Tom^*) mice^32^. To do so, cre-recombination was induced in 8-week-old SC*^Tom^* mice, after which mice were euthanized either seven days post-induction (baseline) or at selected time points up to six months after tamoxifen treatment (Figure 1A). We found that nearly 1% of acinar cells dispersed, throughout the pancreas, expressed tomato at baseline (Figure 1B-D), indicating that these cells actively expressed *Stmn1* at the time of tamoxifen treatment. Next, to determine whether the Stmn1-lineage expands within the acinar compartment during homeostasis, SC*^Tom^* mice were analyzed at 2, 4 or 6 months after tamoxifen treatment (Figure 1B). We found that the number of Tomato-labeled acinar cells increased progressively during this time course to nearly 15% (Fig. 1C). Of note, similar to their 8-week old baseline analysis, we detected tomato expression in approximately 1% of acinar cells in SC*^Tom^* mice that had been treated with tamoxifen at 8 months of age and harvested one week later (Supplementary Figure 1A), suggesting that the percentage of *Stmn1*-expressing acinar cells does not change by age. The constant number of *Stmn1*-expressing acinar cells along with the gradual increase of tomato-labeled acinar cells (Figure 1B, C) implies that the contribution of Stmn1-lineage to acinar tissue homeostasis is likely to be through asymmetric cell division, with an STMN1^+^ cell dividing and giving rise to an STMN1^+^ and an STMN1^-^ cell. We reasoned that the prerequisites for asymmetric division would be that (i) the ratio of Tom^+^/STMN1^+^ cells within the larger tomato-labeled population should decline with time, and (ii) the vast majority (if not all) of STMN1^+^ cells should be tomato-labeled. To test this, tissues obtained from the abovementioned groups of mice were immunostained for STMN1 (Figure 1E & Supplementary Figure 1B). As expected, all Tom^+^ cells in the baseline cohort were STMN1^+^. However, this ratio dropped sharply to 2.5% within the first two months after Cre-activation and continued to near non-detectable levels in the following four months (Figure 1F). Similarly, the percentage of STMN1^+^ cells that were tomato-labeled declined from 93% a week after tamoxifen treatment (reflecting 93% Cre penetration) to almost zero within six months. In contrast, the percentage of Tom^-^/STMN1^+^ increased from 7% (cells that had escaped Cre activity) to essentially 100% during the same period (Figure 1F). Together, these results suggest that maintaining a constant pool of STMN1^+^ cells under physiological conditions and acinar tissue homeostasis relies primarily on recruitment of new *Stmn1*-expressing cells rather than asymmetric cell division within the lineage.

**Figure 1.**
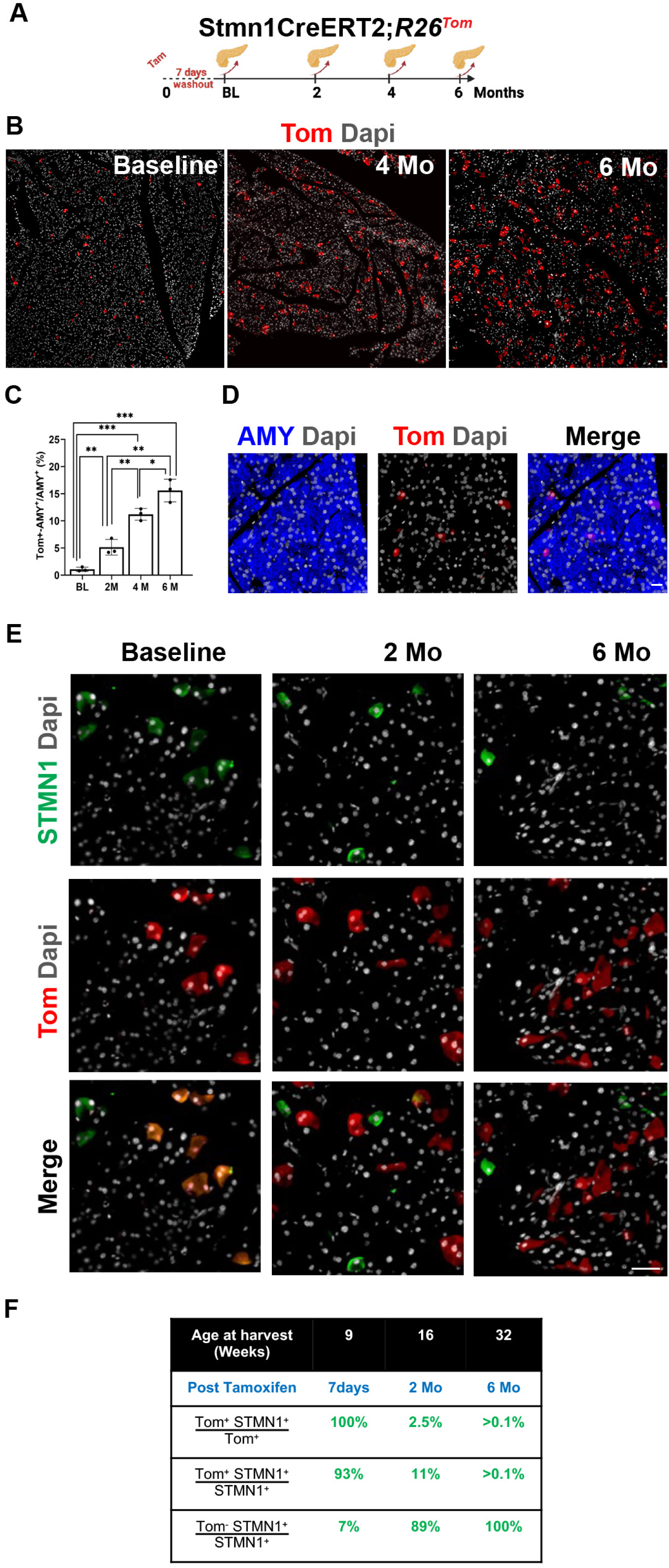
Stmn1-ineage supports acinar homeostasis. (A) Graphical depiction of the lineage tracing strategy. (B, C) Representative data from pancreas sections of SC*^Tom^* mice harvested either 7 days, or 2-, 4-, or 6-months post-tamoxifen treatment (B) and corresponding quantification (C). n = 5; Statistical analysis was performed using unpaired, two-tailed t-test. Data is presented as mean ± SD. * p ≤ 0.05, ** p ≤ 0.01, ***p ≤ 0.001. (D, E) Sections additionally stained for detection of amylase (D) or STMN1 (E). (F) Quantification of the percentage of tomato-labeled cells that actively express *Stmn1* a week, or 2 or 6 months after tamoxifen treatment. Scale bars 20μm. BL: baseline

### Stmn1-lineage plays a pivotal role in acinar regeneration

To identify a role for the STMN1 population during tissue injury in the pancreas, we first interrogated our previous published single cell RNA sequencing (scRNA-seq) dataset generated from caerulein treated pancreata^19^. Caerulein is a cholecystokinin ortholog that induces inflammation, cell death and proliferation, and ADM in the pancreas^38^. To follow the fate of acinar cells upon injury, we performed lineage tracing using the broad, acinar-specific *Ptf1aCre^ERTM/+^;Rosa^LSL-eYFP/+^*(CY) mouse model and induced injury using repeated injection of caerulein. As previously reported, we found that acinar cells are highly plastic in response to injury and form several distinct secretory cell populations during ADM (Figure 2A, Supplementary Figure 2A-D)^19, 39, 40^. As shown in Figure 2B, we identified 2 major *Stmn1*+ populations: one representing *Pcna*+ proliferating acinar cells (Figure 2C) and the other present in ADM. To validate these data, we performed immunostaining on pancreata from (CY) mice treated with tamoxifen, to induce eYFP expression, with or without caerulein treatment. As shown in figure 2D, we identified rare STMN1+, eYFP+ amylase+ acinar cells, consistent with results in SC*^Tom^* mice. Interestingly, we identified PCNA expression in these cells, suggesting that STMN1+ acinar cells represent a small, proliferative population in the normal pancreas. In response to caerulein treatment, we identified STMN1 expression in eYFP^+^/amylase^-^ cells, reflecting ADM. Again, STMN1^+^ cells were often PCNA^+^ consistent with a known role for STMN1 in proliferation^17^ (Figure 2E). To better understand the lineage trajectory of these two populations from normal acinar cells, we performed Pseudotime on our scRNA-seq dataset. Results suggest that *Stmn1*^+^ proliferating acinar cells form earlier in injury than *Stmn1*^+^ ADM populations, further suggesting that ADM populations could be derived from these acinar cells or may represent a second distinct lineage trajectory for acinar cells (Figure 2D). Consistent with our lineage tracing and immunostaining analyses (Figure 1), we identified a small population of *Stmn1*+ acinar cells in normal pancreata, which greatly expand with caerulein-induced injury (Figure 2H-I). Under both conditions, *Stmn1^+^* expression was enriched in *Pcna*^+^/*Mki67*^+^ proliferating acinar cells, as compared to ADM, confirmed by enrichment of S and G2M cell cycle phase scores (Figure 2J, Supplementary Figure 2F-H). To determine if proliferation and ADM represent distinct fates for acinar cells upon injury, we performed Monocle 2 on this modified dataset (Figure 2K). Results suggest that normal acinar cells either become proliferative or undergo ADM. Consistent with Tosti et al., these data suggest the possibility that proliferation and ADM represent distinct cell fates for acinar cells^41^.

**Figure 2.**
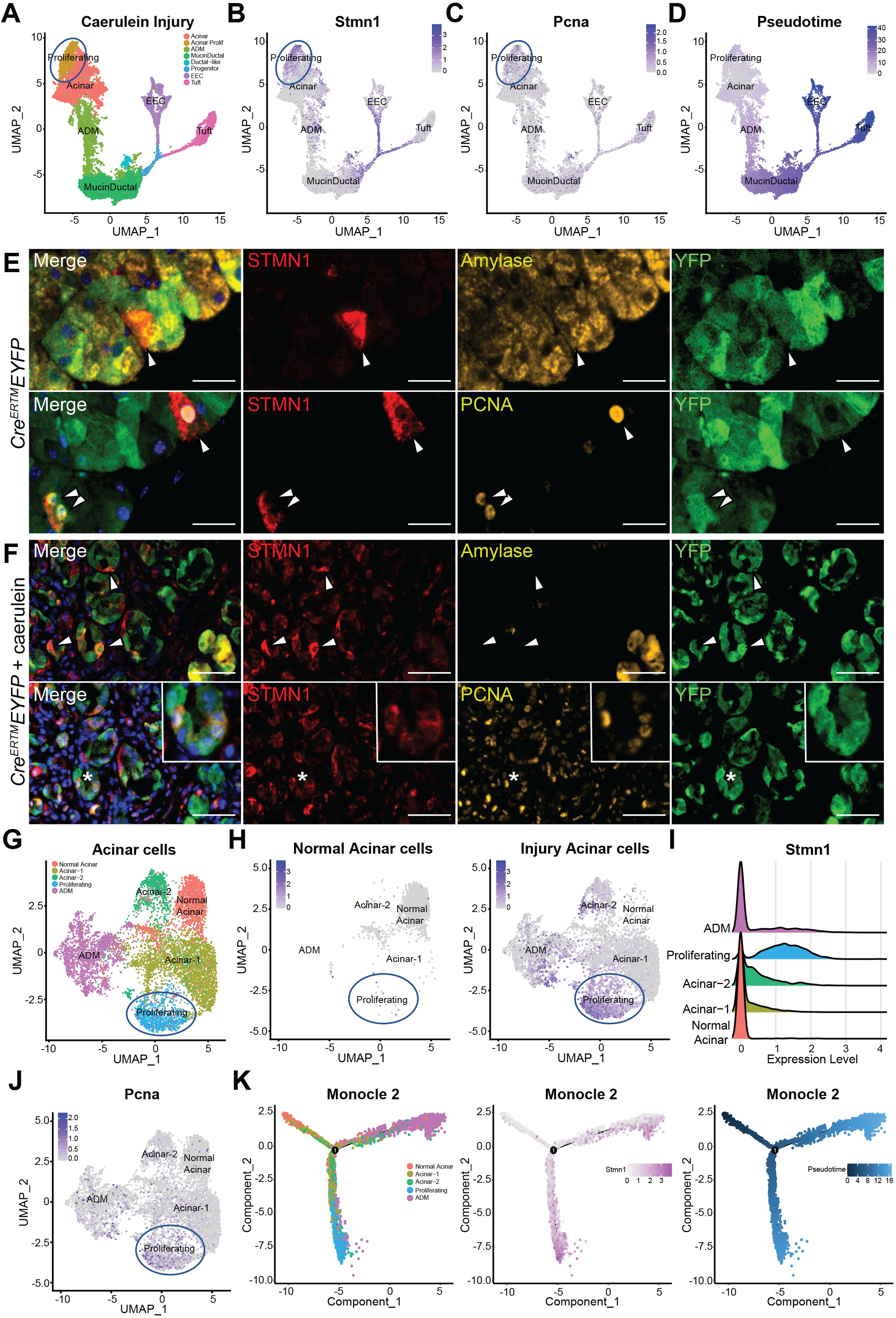
*Stmn1* is expressed in multiple injury-induced populations in the pancreas. (A) Uniform Manifold Approximation and Projection (UMAP) showing annotated clusters in an eYFP+ population derived from *Ptf1aCre^ERTM/+^;Rosa^LSL-EYFP/+^*(CY) mice treated with tamoxifen and caerulein. Expression of (B) *Stmn1* and (C) *Pcna* overlaid on the UMAP of EYFP+ cells. (D) Pseudotime projection analysis on the UMAP from (A). (E) Co-IF for STMN1, amylase or PCNA, YFP and DAPI in CY mice treated with tamoxifen, without injury or in (F) CY mice treated with tamoxifen and caerulein. Scale bars, 25 μm and 50 μm, respectively. (G) UMAP showing select annotated clusters from (A) combined with normal acinar cells from additional scRNA-seq datasets. (H) *Stmn1* expression overlaid on the UMAP in (G) broken out into normal acinar and injury datasets and (I) presented as a Ridge plot. (J) *Pcna* expression overlaid on the UMAP in (G). (K) Monocle 2 trajectory analysis of the populations represented in (G) overlaid with either *Stmn1* expression or pseudotime.

To better understand the role of STMN1^+^ acinar cells in this cell fate decision, we isolated our ‘Acinar’, ‘Proliferating’, and ‘ADM’ populations (Figure 2A) and combined this dataset with scRNA-seq conducted on acinar cells isolated from normal, untreated pancreata (Figure 2G, Supplementary Figure 2)^36, 37^. As shown in supplementary figure 2A&B, acinar cell markers *Cpa1* and *Try4* are highly expressed in normal acinar cell populations but are reduced in ‘ADM’ in which cells are assuming a ductal fate. Consistent with our lineage tracing and immunostaining analyses (Figure 1), we identified a small population of *Stmn1*+ acinar cells in normal pancreata, which appear to greatly expand with caerulein-induced injury (Figure 2G, H). Under both conditions, *Stmn1+* expression was enriched in *Mki67*+, *Pcna*+ proliferating acinar cells, as confirmed by enrichment of S and G2M cell cycle phase scores (Figure 2I, K), to a much greater extent than in ADM. To determine if proliferation and ADM represent distinct fates for acinar cells upon injury, we performed Monacle 2 on this modified dataset (Figure 2L). Results suggest that normal acinar cells either become proliferative or undergo ADM. Collectively these data demonstrate that *Stmn1* is expressed in rare adult acinar cells and greatly expands with injury, first into a proliferative population which does not undergo ADM but instead functions in tissue regeneration and repair.

### Stmn1-lineage proliferates but does not contribute to ADMs

To determine experimentally if the STMN1 population represents a lineage specific for proliferation and acinar regeneration, we used caerulein-induced AP in SC*^Tom^* mice. It is well accepted that this model causes focal injury in some acinar lobes, whereas other lobes are not affected. Interestingly, we observed a significant expansion of Tom^+^ cells specifically in lobes with injury (Figure 3A-D) as evident by the infiltration of macrophages and morphological changes (Supplementary Figure 3A-C). It should be emphasized that the quantification of Tom^+^ cells (Figure 3D) was based only on those lobes with obvious injury (Figure 3C, area marked by dotted yellow line), rather than including uninjured lobes. Transcriptomic analysis on wild type mice have shown that acinar injury leads to an expansion of cells expressing *Stmn1*^17^. Accordingly, and consistent with our computational analyses, we could detect a transient increase in the number of STMN1^+^ cells on day 3 post-caerulein, which declined on day 7 (Figure 3E-F, Supplementary Figure 3D, E). Notably, most of these STMN1^+^ cells were not tomato-labeled (Figure 3G). The presence of Tom^-^/STMN1^+^ cells indicates that some cells express *Stmn1* for the first time after injury. Given the observed new onset of *Stmn1* expression in response to injury, and to distinguish between the original STMN1^+^ cells and those with new onset, we hereafter refer to the original *Stmn1*-expressing cells and their progeny as the Stmn1-lineage.

**Figure 3.**
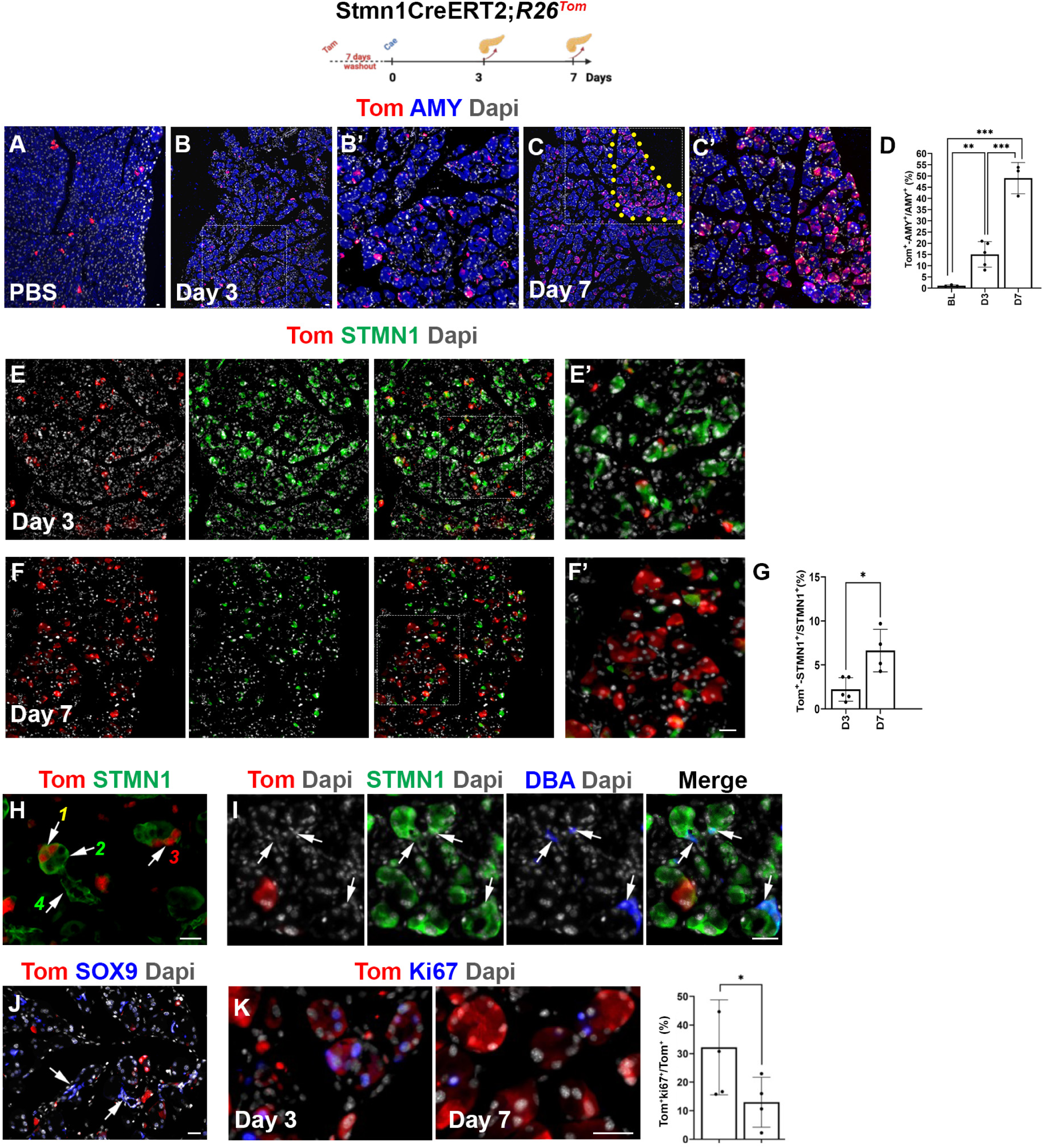
Stmn1-lineage contributes to acinar regeneration in an ADM-independent manner. (A-D) Fluorescent imaging of Tom^+^ cells in SC*^Tom^*pancreas for detection of amylase (A-C) in mice treated with PBS (A) or on days 3 (B) or 7 (C) following caerulein treatment and corresponding quantification (D). (B’, C’) are higher magnifications of the areas highlighted in (B, C). Yellow dotted line in (C) marks a regenerating lobe with the prominent presence of Tom^+^ cells. n = 3; Statistical analysis was performed using unpaired, two-tailed t-test. Data is presented as mean ± SD. ** p ≤ 0.01, ***p ≤ 0.001. (D-G) Fluorescent imaging of Tom^+^ cells in SC*^Tom^* pancreas for detection of STMN1 on days 3 (E) or 7 (F) following caerulein treatment and corresponding quantification (G). n = 3; Statistical analysis was performed using unpaired, two-tailed t-test. Data is presented as mean ± SD. * p ≤ 0.05. (H, I) Fluorescent imaging of tomato-red in conjunction with STMN1 (H) or STMN1 and DBA (I) on day 3 post-caerulein treatment. Arrows in (H) mark Tom^+^/STMN1^+^ acinar cells (*1*), Tom^-^/STMN1^+^ acinar cells (*2*), Tom^-^/STMN1^+^ acinar cells (*3*) or Tom^-^/STMN1^+^ ductular structures (*4*). Arrows in (I) show DBA^+^ ductular structures with no tomato expression. (J) Fluorescent imaging of tomato-red in conjunction with SOX9. Arrows in (J) mark showing the absence of Tom^+^ cells in Sox9-expressing ADMs. (K) Fluorescent imaging of tomato-red and ki67 in the pancreas of SC*^Tom^*mice harvested on days 3 or 7 post-caerulein treatment and the corresponding quantification of the proliferating tomato-labeled acinar cells. n = 3; Statistical analysis was performed using unpaired, two-tailed t-test. Data is presented as mean ± SD. * p ≤ 0.05. Scale bars 20μm.

Given the involvement of ADMs in acinar regeneration and the contribution of the Stmn1-lineage to the regeneration of acinar tissue, we looked for tomato-labeled cells with or without active *Stmn1* expression within ADMs. Our analysis of caerulein-treated SC*^Tom^*mice showed that STMN1 protein was detected in both Tom^+^ (*1* in Figure 3H) and Tom^-^ (*2* in Figure 3H) acinar cells, as well as Tom^-^ ductular structures (Figure *4* in Figure 3H). We also found acinar progeny to the original *Stmn1*- expressing cells (Tom^+^) that were STMN1^-^ (*3* in Figure 3H). The ductular identity of these Tom^-^ structures was further confirmed by their ability to bind lectin-conjugated DBA (Figure 3I). Thus, while the original baseline *Stmn1*-expressing Tom^+^ cells gave rise to both STMN1^-^ and STMN1^+^ acinar progeny, it did not give rise to any ductular structures, including SOX9^+^ ADMs (Figure 3J). Consistent with the expansion of Tom^+^ cells in the context of acinar injury, 30% of Tomato-expressing acinar cells were Ki67^+^ (Figure 3K). These findings indicate that acinar cells within the Stmn1-lineage contribute to acinar regeneration through a process that does not involve ADMs.

**Figure 4.**
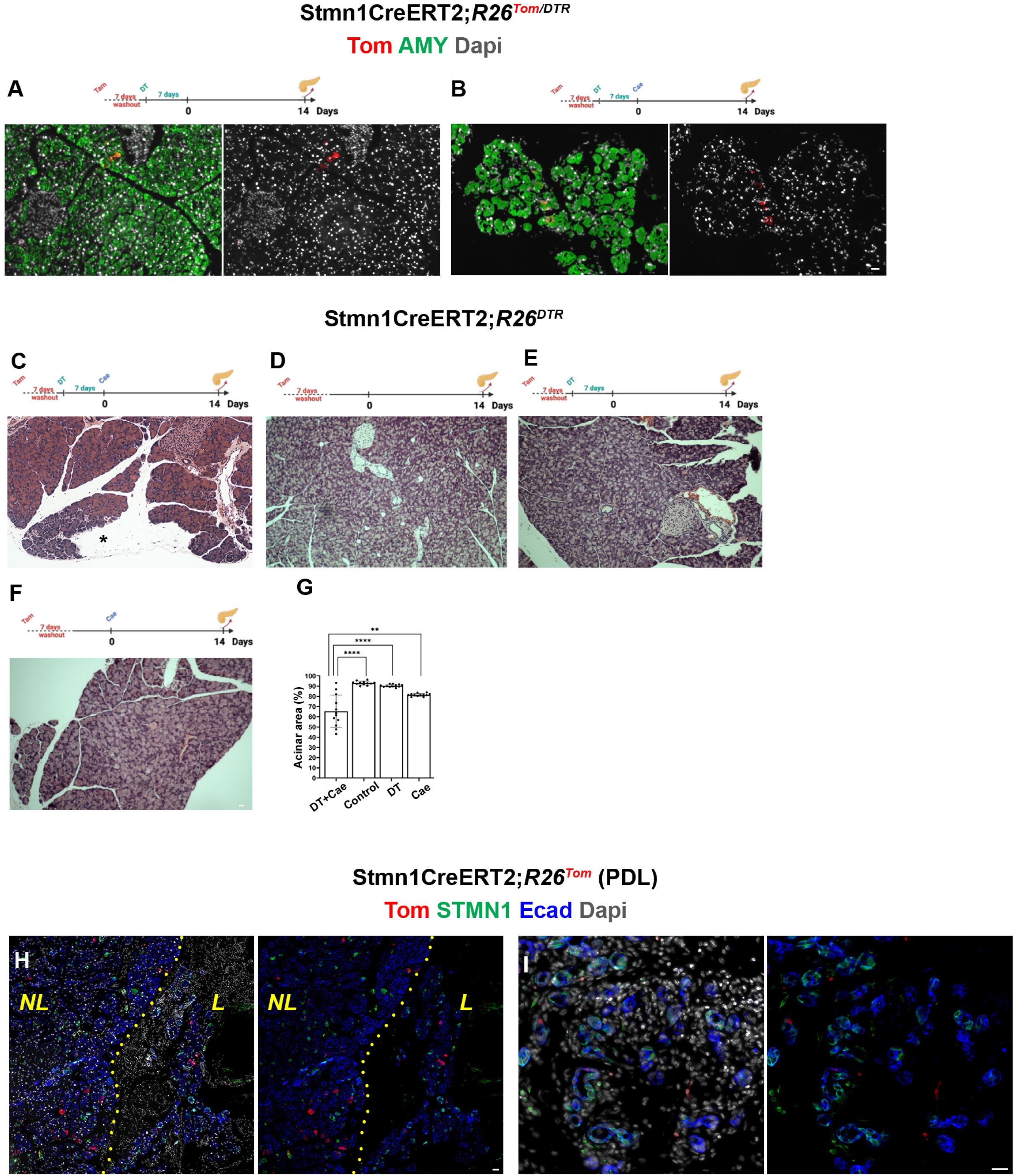
Incomplete acinar regeneration is correlated with loss of the Stmn1-lineage. (A, B) Representative images from pancreas sections of DT-treated SC*^Tom/DTR^* mice harvested without (A) or with caerulein (B) treatment. (C-F) Representative data from pancreas sections of SC*^DTR^*mice treated with DT and caerulein (C), saline (D), DT alone (E) or caerulein alone (F) stained with hematoxylin and eosin. (G) Corresponding quantification of the acinar area shows impaired acinar regeneration in mice treated with DT and caerulein. n = 3; Statistical analysis was performed using unpaired, two-tailed t-test. Data is presented as mean ± SD. ** p ≤ 0.01, ****p ≤ 0.0001. Scale bars 20μm. (H, I) Fluorescent imaging of Tomato in conjunction with STMN1 and E-cadherin in the pancreas of SC*^Tom^* mice harvested 2 weeks post PDL. Note the absence of Tomato-expressing cells in the ligated part. *NL*: Ligated; *L*: Ligated. Scale bars 20μm.

### Incomplete acinar regeneration is correlated with the loss of the Stmn1-lineage

To better assess the importance of *Stmn1*-expressing cells to acinar regeneration, we generated Stmn1CreERT2;*R26^Tom^*^/*DTR*^ (*SC^Tom/DTR^*) mice in which the diphtheria receptor (DTR) expressing cells will undergo an apoptotic death upon exposure to diphtheria toxin (DT). This cell ablation approach has been previously used by our lab to study pancreatic regeneration in the context of cell-specific ablation of different cell types in the adult pancreas^35, 42, 43^. Here, administration of DT to *SC^Tom/DTR^* mice prior to injury would ablate the Stmn1-lineage, but not cells with the new onset of *Stmn1* expression after injury, thus allowing us to study acinar regeneration in the absence of the original *Stmn1*-expressing cells. We thus administered DT to SC*^Tom/DTR^*mice and divided them in two cohorts; one cohort was left alone (Figure 4A & Supplementary Figure 4A), whereas the other group was treated with caerulein (Figure 4B & Supplementary Figure 4B). Two weeks later, the pancreas of DT-treated mice looked indistinguishable from a normal control pancreas, whereas the pancreas of mice treated with DT and caerulein showed impaired acinar regeneration (Figure 4A, B & Supplementary Figure 4A, B). Furthermore, in both cohorts we could hardly find any Tom^+^ cells (Figure 4A, B & Supplementary Figure 4A, B), confirming DT/DTR-mediated ablation of the Stmn1-lineage.

To better quantify acinar regeneration in the absence of the Stmn1-lineage, we used tamoxifen-treated Stmn1CreERT2;*R26^DTR^* (*SC^DTR^*) mice. These mice were then divided in six cohorts based on whether they received DT+caerulein (Figure 4C), no additional treatment (Figure 4D), only DT (Figure 4E) or only caerulein (Figure 4F) and harvested on day 3 (Supplementary Figure 4C, D) or 14 post caerulein treatment (Figure 4C-D). Two weeks after caerulein treatment, the pancreas of (DT+caerulein)-treated SC*^DTR^* mice displayed 60% acinar content, whereas the pancreas of SC*^DTR^* mice treated only with caerulein (no DT) showed 80% acinar content (Figure 4G). SC*^DTR^* mice treated only with DT had similar acinar content as the control mice (Figure 4G). Of note, the pancreas of caerulein-treated *SC^DTR^* mice on day 3 post-caerulein displayed a similar extent of initial acinar injury regardless of DT treatment (Supplementary Figure 4C, D). Interestingly, acinar lobes were partially missing in the (DT+caerulein)-treated SC*^DTR^*mice (Figure 4C), reflecting the focal nature of this injury model (Supplementary Figure 4C, D). This result further supports the importance of the Stmn1-lineage in acinar regeneration.

To further confirm the reliance of acinar regeneration on the Stmn1-lineage and to rule out any model specific effect, we performed pancreatic duct ligation (PDL) in SC*^Tom^* mice. In mice, although PDL results in the formation of ADMs, the acinar compartment does not regenerate ^11, 44–46^. Unlike caerulein-mediated injury, where we observed cells with new onset of *Stmn1* expression (Tom^-^/STMN1^+^) only in the injured areas, in the two weeks post-PDL pancreas, Tom^-^/STMN1^+^ cells could be detected in both the ligated part and the surrounding non-ligated tissue (Figure 4H). As expected, PDL resulted in formation of ductular structures (Figure 4H, I), which have been identified as either ADMs^11^ or pre-existing ducts that have changed morphology^44^. Interestingly, some of these structures expressed STMN1 and others did not (Figure 4I). Based on our previous findings, we believe ductular structures with new onset of *Stmn1* expression are ADMs, whereas the STMN1-negative structures are preexisting ducts. More importantly, we did not find an expansion of the Stmn1-lineage in the ligated part (Figure 4H, I). Together, these data show that in the absence of the Stmn1-lineage, acinar regeneration is impaired.

### Stmn1-lineage during CP progression

It has been reported that a shorter caerulein treatment (4 weeks) is correlated with regeneration of the acinar mass (formation of new acinar cells) to the near pre-injury levels, whereas recovery after a longer period of insult (10 weeks) primarily entails repair (no new acinar cells)^47, 48^. Based on the key role the Stmn1-lineage plays in acinar regeneration following caerulein-induced AP, we next studied whether the reported switch between regeneration and repair would be associated with a decline in the Stmn1-lineage during CP progression. As a first step, wild type mice were treated with caerulein for either four weeks (Supplementary Figure 5A) or eight weeks (Supplementary Figure 5B). In the 4-weeks cohort, we could find STMN1^+^ cells scattered within acinar compartment and in ADMs (Supplementary Figure 5C). However, STMN1 detection in the 8-weeks cohort was restricted to ADMs and acinar cells seemingly undergoing ductal metaplasia (Supplementary Figure 5D, E). Since this study was performed in wild type mice, we could not distinguish between the Stmn1-lineage and cells with new onset of *Stmn1* expression. To address this question, following tamoxifen treatment, SC*^Tom^* mice received caerulein for four or eight weeks, after which the mice were either euthanized (Figure 5A & Supplementary Figure 6A, C) or were allowed to recover for four weeks (Figure 5A & Supplementary Figure 6B, D), while mice treated with PBS served as controls (Figure 5B). As expected, the total pancreatic area consisted of approximately 95% amylase^+^ acinar cells in the control pancreas, and Sirius red stained only the area only surrounding the larger ducts (Figure 5B, G). Four weeks of caerulein treatment resulted in 70% reduction in acinar area and minor presence of fibrosis (Figure 5C, G), whereas the subsequent four weeks of recovery lead to a 1.6x increase in acinar area and regaining 50% of the original acinar content (Figure 5D, G). As shown in figure 5, we observed near 80% loss of acinar tissue and higher degree of fibrosis in mice that received eight weeks of caerulein treatment (Figure 5E, G). However, unlike the four-weeks regiment, allowing mice to recover following eight weeks of caerulein treatment did not result in an increase in acinar area (Figure 5F, G). Of note, the partial regeneration in the (4w cae+4w rec) cohort was in conjunction with a higher ratio of Tom^+^/amylase^+^ cells compared to the (4w cae) group (Figure 5C, D, H). However, we could not detect any increase in the Tom^+^/amylase^+^ ratio in the (8w cae+4w rec) cohort compared to the (8w cae) mice (Figure 5E, F, H). Perhaps more importantly, we could hardly detect any Tom^+^/STMN1^+^ cells in the pancreas of (8w cae+4w rec) mice (Figure 5F, I). This is highly relevant, as cells with active *Stmn1* expression within this lineage (Tom^+^/STMN1^+^ cells), are likely to be responsible for proliferation and lineage maintenance. Finally, the clonal expansion of Tom^+^ cells observed in the (4w cae+4w rec) cohort was absent in the (8w cae+4w rec) pancreas (Figure 5D, F). Collectively, these data suggest that expansion of the Stmn1-lineage which occurs under regenerative conditions (AP or early stages of CP) is either not observed or heavily impaired under conditions leading to repair.

**Figure 5.**
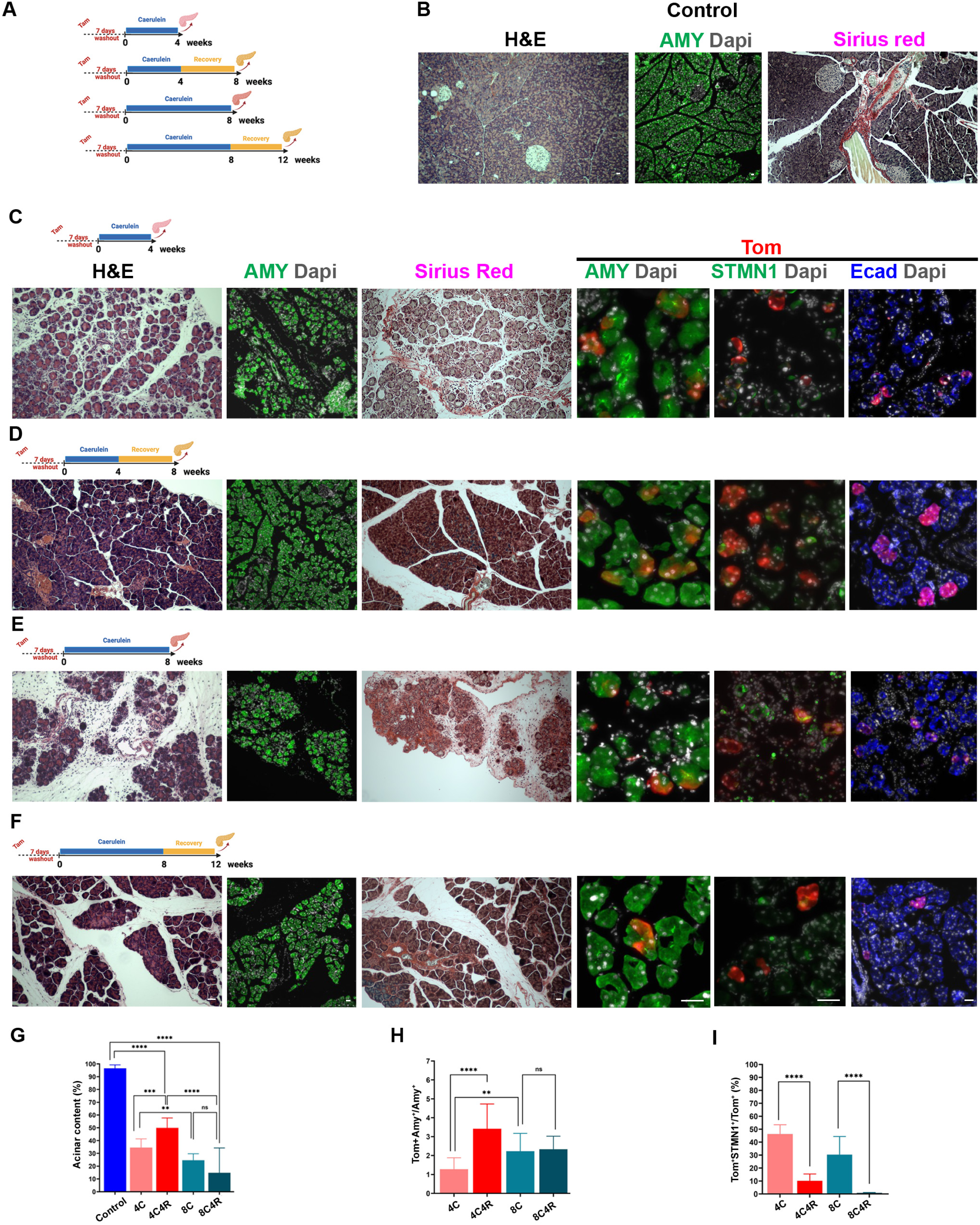
Acinar regeneration during CP progression is correlated with the proliferative capacity of the Stmn1-lineage. (A) Graphical illustration of the chronic pancreatitis cohorts. (B) Sections from the control pancreas stained with hematoxylin and eosin, Sirius red or stained for detection of amylase. (C-F) Sections stained with hematoxylin and eosin, Sirius red or stained for detection of tomato-red in conjunction with amylase, STMN1 or E-cadherin from the pancreas of SC*^Tom^* mice harvested after 4 weeks caerulein treatment (C), 4 weeks caerulein treatment and 4 weeks recovery (D), 8 weeks caerulein treatment (E), or 8 weeks caerulein treatment and 4 weeks recovery (F). (G-I) Graphs show quantification of the regenerated acinar compartment (G), percentage of Tom^+^ acinar cells (H), or percentage Tom^+^ acinar cells with active *Stmn1* expression in the abovementioned cohorts. n = 3; Statistical analysis was performed using unpaired, two-tailed t-test. Data is presented as mean ± SD. ^ns^ p > 0.05, * p ≤ 0.05, ** p ≤ 0.01, ***p ≤ 0.001, ****p ≤ 0.0001. Scale bars 20μm.

### Stmn1-lineage does not contribute to pancreatic neoplasia upon oncogenic Kras expression

ADM is considered as to be a transition state between the acinar cells and PanINs in the context of oncogenic *Kras* expression in mouse models of pancreatic cancer^13, 15, 49^. However, the pan-pancreatic expression (using Pdx1Cre or Ptf1aCre mice) or pan-acinar expression (using ElaCreERT, Mist1CreERT or Ptf1aCreERT mice) of oncogenic Kras in these mouse models have been a limitation to determine whether ADM is an absolute prerequisite for PanIN formation. Given our finding that STMN1-lineage acinar cells contribute to tissue regeneration but do not undergo ADM in response to injury, we hypothesized that this population of lineage restricted acinar cells would not contribute to neoplasia and tumor formation. To first assay for *Stmn1* expression in pancreatic neoplasia, we compared our injury scRNA-seq dataset to one generated from *Kras^G12D^*-expressing mice by Schlesinger et al^36^. Here, the authors performed lineage tracing of adult acinar cells using the *LSL-Kras^G12D^;Ptf1aCre^ERTM^;LSL-tdTomato* (KCT) model. *Kras^G12D^* and *tdTomato* expression was induced with tamoxifen and pancreata were collected at several timepoints encompassing early changes in the pancreas as well as tumor formation (Figure 6A). Interestingly, we identified few proliferating acinar cells in the *Kras* dataset. Instead, *Stmn1* expression was largely enriched in a smaller population of proliferating ADM, which was not identified in our injury dataset (Figure 6B). Proliferation in both acinar cells from the injury dataset and *Kras^G12D^*-induced ADM was confirmed by expression of *Pcna*, *Mki67,* and several cell cycle regulators (Figure 6C-D). To confirm *Stmn1* expression in proliferative PanIN populations, we performed immunostaining on pancreata from *LSL-Kras^G12D^;Ptf1aCre^ERTM^;Rosa^LSL-EYFP/+^* (*KCYFP*) mice treated with tamoxifen and a short course of caerulein to accelerate disease progression. As shown in Figure 6E, PCNA^+^/STMN1^+^ populations were abundant in acinar-derived PanIN, likely representing the ‘proliferating ADM’ population.

**Figure 6.**
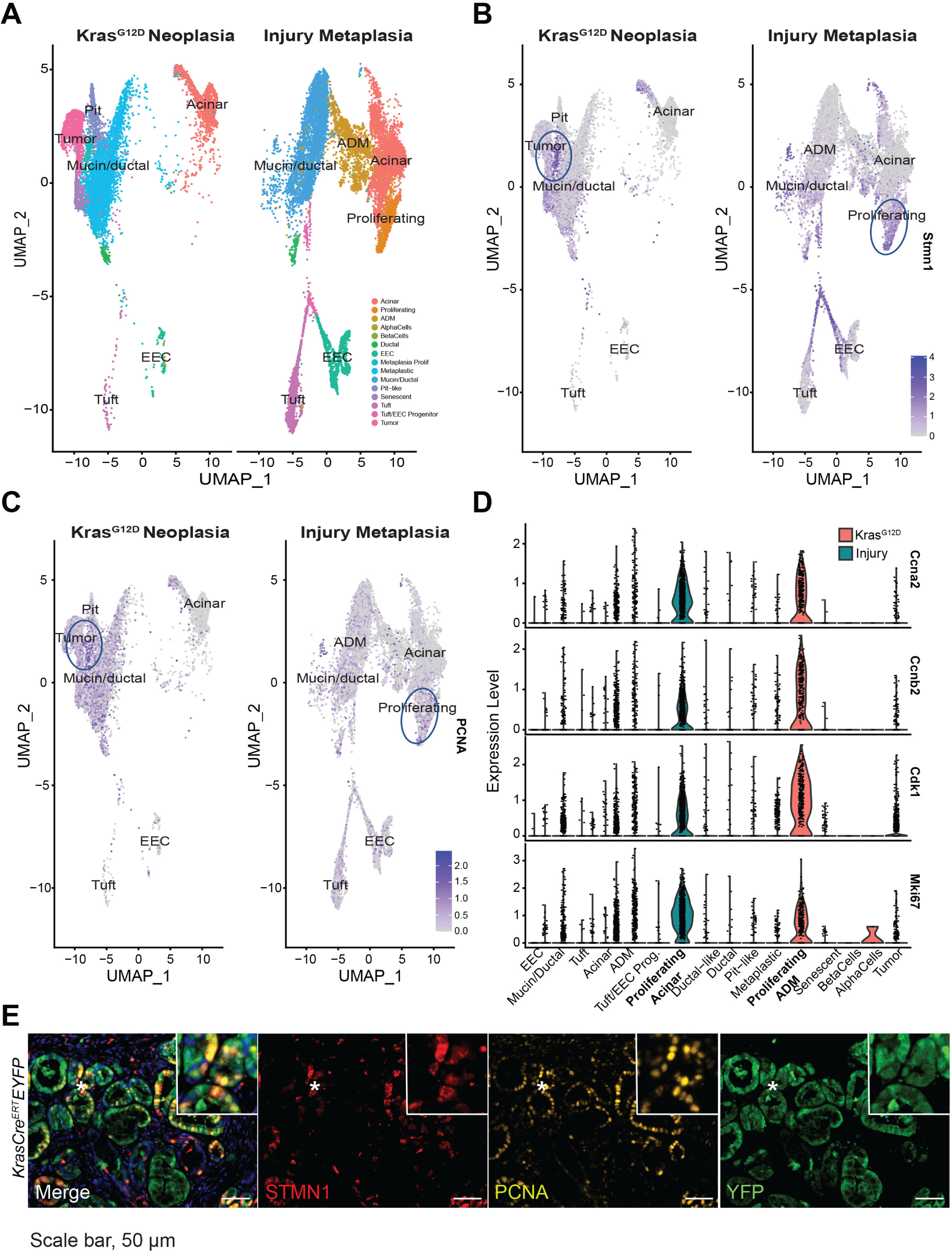
*Stmn1* is expressed in acinar-derived PanIN populations. (A) UMAP of FastMNN integrated datasets from the injury-induced ADM scRNAseq dataset a *Kras^G12D^*-induced dataset of neoplasia and cancer. Expression of (B) *Stmn1* or (C) *Pcna* overlaid on the integrated UMAP. (D) Violin plots of several cell cycle genes and *Mki67* in both datasets. (E) Co-IF for STMN1 (red), PCNA (yellow), YFP (green), and DAPI in PanIN derived from a *KCYFP* mouse treated with tamoxifen and caerulein. Scale bars, 50 μm.

To determine whether the Stmn1-lineage could form neoplastic lesions and contribute to the proliferating ADM population upon *Kras^G12D^*expression despite its inability to contribute to ADMs, we generated Stmn1CreERT2; R26*^Kras^* (SC*^Kras^*) mice. The pancreas of SC*^Kras^* mice treated with corn oil or tamoixifen alone looked normal and were indistinguishable (Figure 7A, B). This observation could be due to the relatively low number of *Stmn1*-expressing cells under physiological conditions, as shown in figure 1 (>1% of total acinar cells in the adult pancreas). One round of caerulein-induced acinar injury leads to expansion of Stmn1-lineage and its contribution to acinar regeneration (Figure 3A-C). In addition, this treatment has been widely used to accelerate PanIN formation and progression^50^. However, in tamoxifen-treated SC*^Kras^*mice subjected to one round of caerulein, no lesions could be found at neither four- nor twelve-weeks post-caerulein treatment (Figure 7C). Here, while caerulein-treatment might have increased the number of Stmn1-lineage-derived Kras-expressing acinar cells, our inability to detect PanINs could be due to the withdrawal of immune cells, as generation of the Stmn1-lineage-derived new acinar cells coincides with the resolution of injury and inflammation. To address this issue, we next subjected tamoxifen-treated SC*^Kras^* mice to two consecutive rounds of caerulein treatments and harvested the pancreas one or six months after the cessation of caerulein treatment (Figure 7D). The pancreas of one-month post-caerulein cohort contained a few lobes with prominent acinar atrophy. Interestingly, no such phenotype could be found in the six-months cohort (Figure 7D). More importantly, similar to previous conditions, we could not find PanINs following two rounds of caerulein treatments. Since injury promotes the new onset of *Stmn1* expression among acinar cells (Figure 3D), to confirm oncogenic *Kras* expression in our mouse model, SC*^Kras^* mice were treated first with caerulein and then with tamoxifen (Supplementary Figure 7). We reasoned that by doing so we would target *Kras* expression not only to Stmn1-lineage, but also to acinar cells with new onset of *Stmn1* expression as well as ADMs. Here, we could observe PanINs as early as four weeks post-tamoxifen treatment (Supplementary Figure 7), suggesting that the absence of lesions when acinar injury was induced after Cre-recombination (Figure 7C, D) is unlikely to be due to suboptimal *Kras* expression.

**Figure 7.**
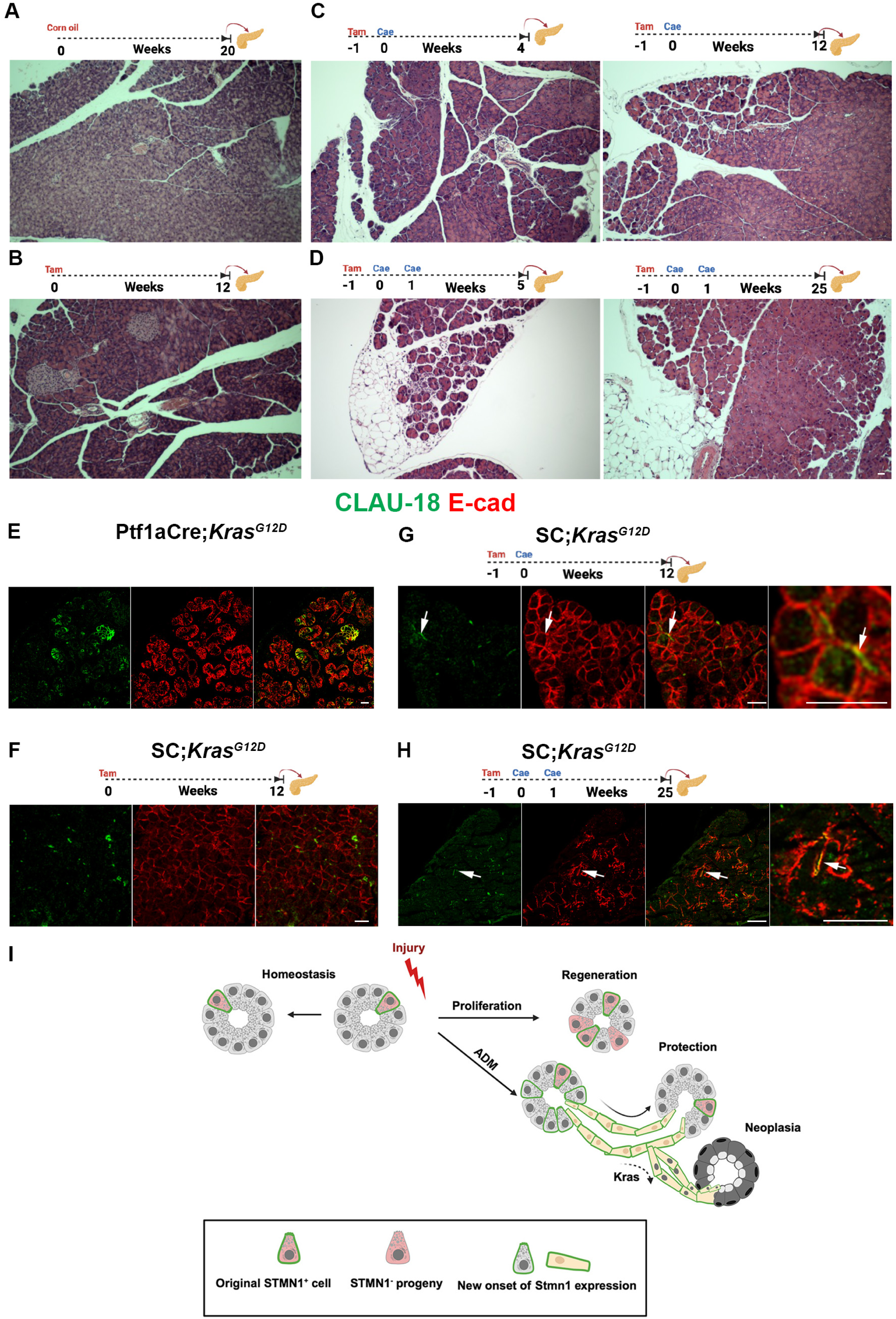
Stmn1-lineage is refractory to Kras*^G12D^* expression. (A-D) Representative images from pancreas sections of SC*^Kras^* mice harvested 20 weeks after corn oil treatment (A), harvested 12 weeks post-tamoxifen treatment (B), treated with tamoxifen first and harvested 4 or 12 weeks post-caerulein treatment (C), or treated with tamoxifen first and harvested 5 or 25 weeks after two rounds of caerulein treatments (D). (E-H) Confocal fluorescent imaging of pancreas sections of Ptf1aCre;*Kras^G12D^* (E), or SC*^Kras^* mice harvested12 weeks post-tamoxifen treatment (F), treated with tamoxifen first and harvested 12 weeks post-caerulein treatment (G), or treated with tamoxifen first and harvested 25 weeks after two rounds of caerulein treatments (H) for detection of caudin-18 and E-cadherin. Arrows in (G, H) highlight colocalization of claudin-18 and E-cadherin in some acinar cells. (I) Illustration of the proposed model for tissue homeostasis, regeneration and neoplasia. Scale bars 20μm.

PanINs are neoplastic structures with distinct morphological features. To look for more subtle molecular changes, we studied expression of claudin-18 (Clau-18) in SC*^Kras^*mice. Clau-18 is an early marker for pancreatic carcinogenesis, as it is excluded from acinar cells, duct cells and ADMs but can be found in the earliest PanINs^51^ (Figure 7E). Interestingly, in caerulein-treated SC*^Kras^* mice, we could find morphologically normal acinar cells that expressed Clau-18 (Figure 7F-H). Together, our findings show that Stmn1-lineage acinar cells are unable to form PanINs upon *Kras^G12D^* expression. Furthermore, these results provide direct evidence that ADM is a necessary transition state between acinar cells and PanINs.

## Discussion

Although, the notion of acinar cell heterogeneity is widely accepted, reports differ on the extent to which various subpopulations may contribute to tissue homeostasis, regeneration or neoplasia^17–23, 41^. These discrepancies have given rise to two main schools of thought on this subject; one supporting the presence of specialized cells, which are normally responsible for homeostasis and regeneration, but could give rise to neoplastic lesions in the context of oncogenic *Kras* expression^18, 21–23^. Another opposes the existence of such intrinsic cellular hierarchy and propose that all acinar cells could be equally involved in these processes^20^. The current results complement previous studies by favoring the presence of such specialized cells during regeneration and neoplasia but not tissue homeostasis.

In this study, we demonstrate that similar to cells expressing *Bmi1*, *Dclk1*, *Tert* or *Tff2*^18, 21–23^, *Stmn1*-expressing acinar cells constitute a small but stable population that contribute to tissue homeostasis under baseline conditions. The involvement of Bmi1-, Dclk1-, Tert-, Tff2- and Stmn1- lineages in tissue maintenance under normal physiological conditions indicates that acinar tissue homeostasis is likely not dedicated to a specific lineage but instead may require equal contribution from all acinar cells regardless of lineage. Notably, what separates the Stmn1-lineage in this regard is the notion that the gradual increase in the lineage-labeled (Tom^+^) acinar cells over time during homeostasis is associated with a decline in Tom^+^ cells actively expressing *Stmn1* to a negligible ratio. This finding suggests that lineage maintenance is likely through recruitment of new *Stmn1*-expressing cells rather than cell division. The appearance of acinar cells with new onset of *Stmn1* expression is not specific for tissue homeostasis. Upon caerulein-induced acute injury, we observed a significant expansion of the Stmn1-lineage, prominently in injured lobes, indicating that this lineage expands in response to injury and contributes to acinar regeneration. Interestingly, we could also detect new onset of *Stmn1*- expression, as evident by the appearance of STMN1^+^/Tom^-^ acinar cells and ADMs. This data shows that the overall expansion of *Stmn1*-expressing cells during acinar regeneration, as previously reported^17^, is due to expansion of the Stmn1-lineage (cells originally expressing *Stmn1*) as well as cells expressing *Stmn1* for the first time.

To assess the requirement of the Stmn1-lineage in regeneration, we next studied acinar regeneration in the absence of the Stmn1-lineage by utilizing the DT/DTR-mediated cell ablation. We found that the regenerative ability in *SC^DTR^* mice treated with DT and caeruelin was significantly reduced compared to mice treated with DT or caerulein alone. Accordingly, we found near absence of the Stmn1-lineage following pancreatic duct ligation, a condition associated with diminished acinar regeneration in mice^11, 44–46^. Collectively, this data shows that incomplete acinar regeneration in the abovementioned injury models is associated with a significant reduction in the Stmn1-lineage, further reinforcing the pivotal role that the Stmn1-lineage plays in acinar regeneration.

In a well-established animal model for CP, repeated injections of caerulein over several weeks causes acinar atrophy, collagen deposition and fibrosis. In this model, while it has been demonstrated that cessation of caerulein treatment leads to regression/reduction in inflammation and fibrosis^47, 48, 52, 53^, there are conflicting reports on whether the acinar compartment is only repaired (healing without regaining its pre-injury mass, i.e. no new acinar cells) or undergoes regeneration (regaining its pre-injury mass, with formation of new acinar cells) during the recovery period^47, 48^. We reasoned that this inconsistency may reflect the differences in caerulein regimens, in particular the duration of treatment. Based on the importance of the Stmn1-lineage in acinar regeneration during AP, we hypothesized that the choice between acinar regeneration (formation of new acinar cells) and repair (no new acinar cells) may be dictated by the viability and proliferative ability of the Stmn1-lineage. Our studies confirmed the inverse relationship between the duration of injury and the regenerative ability of the acinar compartment. Moreover, the absence of STMN1^+^/Tom^+^ acinar cells following prolonged injury suggest that the Stmn1-lineage is depleted and/or its proliferative capacity is diminished under chronic inflammatory conditions, shifting the pancreas from a regenerative to a reparative state. However, their decline during CP progression should be considered as a limitation under chronic stress. Future studies should unravel therapeutic approaches to sustain or restore the Stmn1-lineage in chronic disease settings, with the goal of promoting pancreatic regeneration and preventing irreversible tissue loss.

It is well accepted that multiple episodes of AP may lead to CP^3^. The overall risk of recurrent attacks after the first episode of AP is around 20%, among which approximately 20-25% will eventually develop CP^54–60^. The reason why 25% of all recurrent AP (RAP) cases will develop into CP is not well defined, as RAP is associated with multiple etiologies, clinical variables, and outcomes^54^. As demonstrated here, in the lobes that had been affected by the caerulein-induced AP, we could find up to 50% contribution of Tom^+^ cells on day 7. Of note, following 4 weeks of caerulein treatment (CP model), we could not find lobes with such a prominent presence of Tom^+^ cells. Instead, Tom^+^ cells were often found as either single cells or few adjacent cells scattered throughout the acinar compartment. This finding indicates that acinar cells which were generated after the first round of caerulein treatment and thus should have contributed significantly to the acinar mass in the affected lobes, were likely to be more susceptible to the additional insults than the preexisting acinar cells, and succumbed. The ability of the human acinar compartment to regenerate in disease conditions such as pancreatitis is yet to be proven. Our finding on the potential increased susceptibility of the newly generated acinar cells to injury suggests that the human acinar cells may have the ability to regenerate but perhaps have less chance to survive the additional insults. As RAP lies on the spectrum between AP and CP, further studies are necessary to determine whether it is the time interval between each attack rather than the number of attacks that plays a decisive role in the transition/progression from RAP to CP.

Injury and inflammation induce formation of ADMs, which are considered to play an active role in acinar regeneration^13^. Given the contribution of the Stmn1-lineage to acinar regeneration, it was surprising that we did not detect any Tomato-labeled cells in any DBA^+^ ductular structures, including SOX9^+^ ADMs. Our finding challenges the dogma that these metaplastic structures are actively involved in acinar regeneration, and further supports a previous report showing that the primary purpose of ADMs is to limit tissue damage via a rapid decline in zymogen production^61^ or suppressing inflammation by harboring tuft cells in the context of Kras*^G12D^* expression^62^. Compared to *Bmi1^+^*, *Dclk1^+^* or *Tert^+^*cells^18, 21, 22^, our results highlight the higher contribution of the Stmn1-lineage to acinar regeneration during acute injury. However, unlike these facultative cells, which can contribute to neoplasia, the Stmn1-lineage fails to initiate such a process. The inability of the Stmn1-lineage to undergo acinar-to-ductal metaplasia may also explain why they fail to form PanINs in the context of oncogenic *Kras* expression.

In conclusion, the current study offers a new paradigm for acinar heterogeneity in the context of tissue homeostasis, regeneration and neoplasia (Figure 7I). According to this model, tissue homeostasis relies on the equal contribution of all acinar cells regardless of lineage. Furthermore, successful acinar regeneration entails two concurrent events, one repopulating the pancreas with new acinar cells and the other protecting the surviving cells from further damage. In that regard, the Stmn1-lineage expands and supports generation of new acinar cells. Meanwhile, cells expressing *Bmi1*, *Dclk1*, or *Tert* are more likely to be involved in protecting the organ by responding to the environmental cues, acquiring progenitor-like features (such as new onset of *Stmn1* expression) and subsequently forming ADMs. Given their ability to form ADMs, the latter acinar populations can then initiate neoplasia upon oncogenic *Kras* expression. Our findings along with previous studies oppose the overarching assumption that the same subset of acinar cells that are responsible for tissue maintenance following injury are likely to act as the cells of origin for neoplasia. Our data suggest that pancreatic neoplasia should be defined as a protection mechanism highjacked by oncogenic KRAS.

## Acknowledgments

This work was supported by NIH grants R01DK120698 (to GKG and F.E.), R21AI158824 (to F.E), CA236965 (to F.E), 1K08DK129834 (to M.S), JDRF grant 2-SRA-2022-1211-S-B (to F.E.), Research Advisory Committee (RAC) Children’s Hospital of Pittsburgh of UPMC (to F.E), Cochrane-Weber Endowed Funds for Diabetes Research (to F.E.) and The Children’s Hospital of Pittsburgh of UPMC (to F.E.). The DelGiorno laboratory was supported by the Vanderbilt Ingram Cancer Center Support Grant (NIH/NCI P30CA068485), the Vanderbilt-Ingram Cancer Center SPORE in Gastrointestinal Cancer (NIH/NCI P50CA236733), the Vanderbilt Digestive Disease Research Center (NIH/NIDDK P30DK058404), an American Gastroenterological Association Research Scholar Award (AGA2021-13), NIH/NIGMS R35GM142709, The Department of Defense (DOD W81XWH2211121), The American Cancer Society (CAT-24-1374680-01-CAT), The Sky Foundation, Inc (AWD00000079), and Linda’s Hope (Nashville, TN). The Leica Stellaris 5 confocal microscope at the Rangos Research center Cell Imaging core facility was purchased using an internal support grant from the Children’s Hospital. We acknowledge the Cell imaging core staff for their support.

## Author contributions

K.E.D and F.E designed the experiments. S.D., and J.R.A. performed the lineage tracing studies under physiological conditions or after caerulein-induced acute pancreatitis. S.D., J.R.A., A.H. S.A., M.W and K.G performed the lineage tracing studies after caerulein-induced chronic pancreatitis. S.D. generated bone marrow-derived macrophages and performed studies involving pancreatic tissue slices. M.S. performed the pancreatic duct ligation as well as chronic pancreatitis in wild type mice. B.K.K. generated the Stmn1CreERT strain. S.D., J.R.A. and F.E performed and analyzed the Sc*^Kras^* studies. Z.M., C. C., and K.E.D performed the bioinformatic analysis. S.D., K.E.D. and F.E interpreted results. K.E.D and F.E. wrote the original draft. K.E.D., G.K.G., and F.E. reviewed and edited the manuscript.

## Supplementary Figure Legends

**Supplementary Figure 1.**
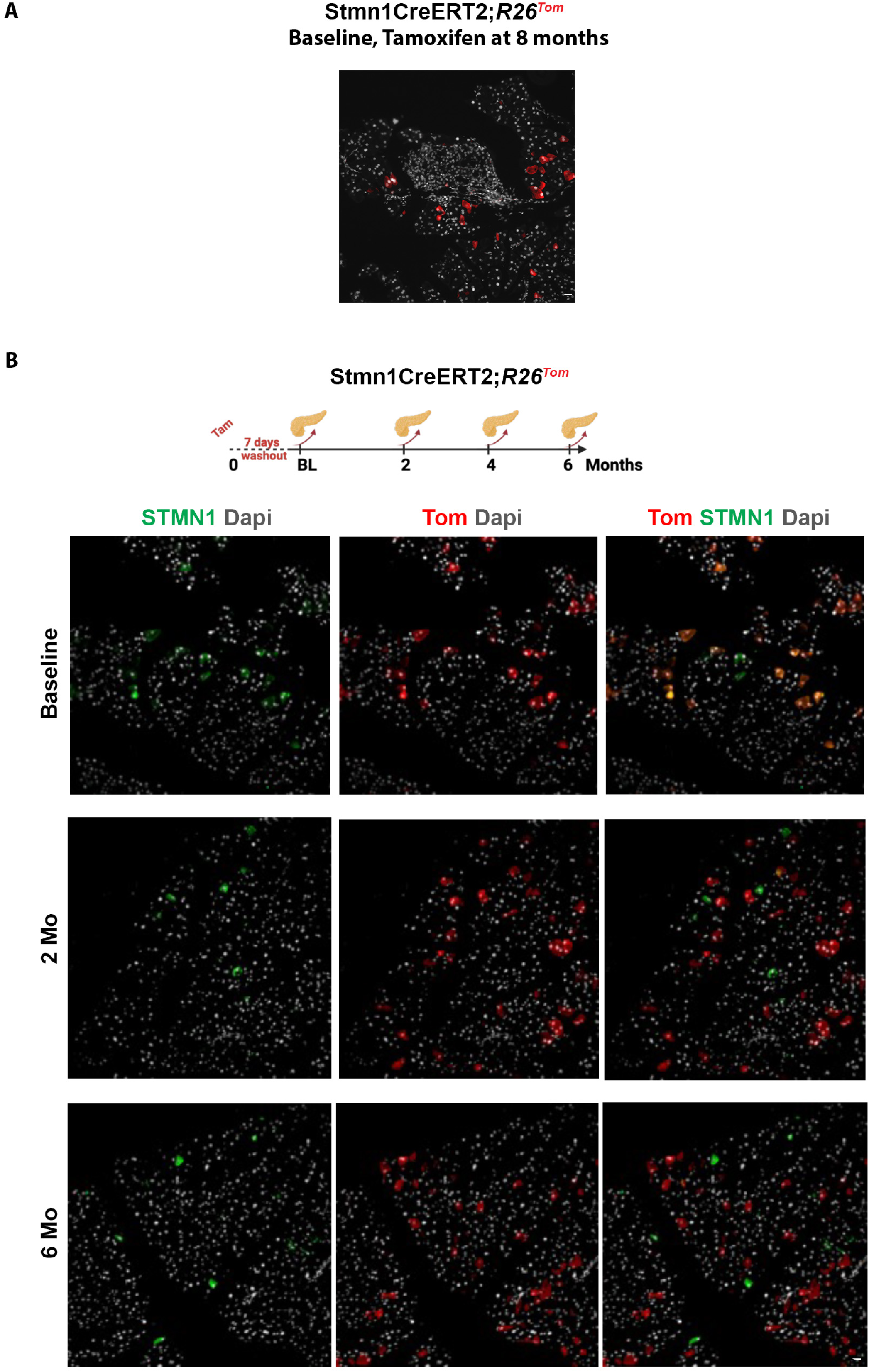
Stmn1-ineage supports acinar homeostasis. (A) Fluorescent imaging of Tom^+^ cells in 8 months old SC*^Tom^* pancreas harvested 7 days post-tamoxifen treatment. (B) Representative data from pancreas sections of SC*^Tom^* mice harvested either 7 days, or 2-, or 6-months post-tamoxifen treatment stained for detection of amylase STMN1. Scale bars 20μm.

**Supplementary Figure 2.**
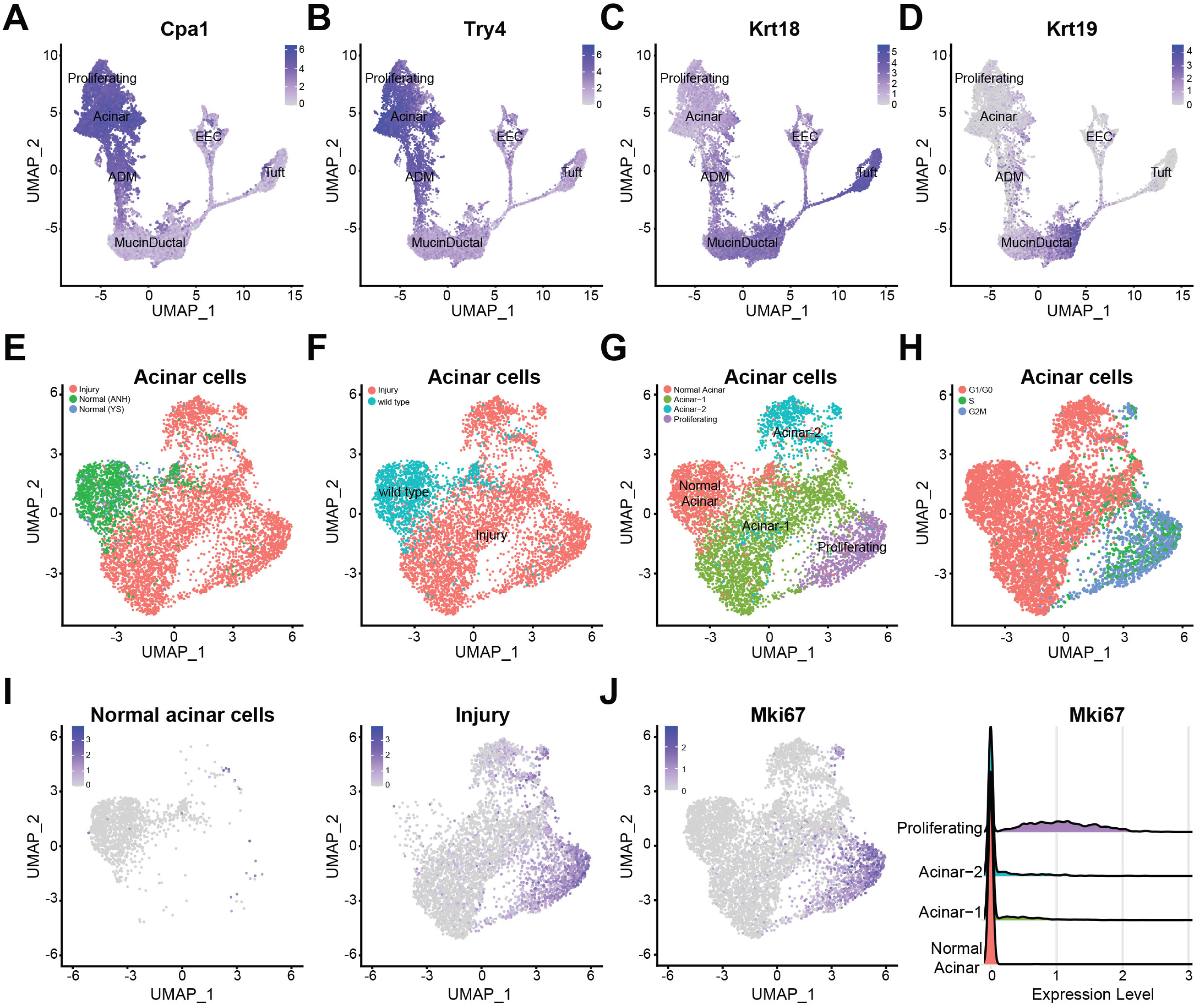
*Stmn1* is expressed in proliferating acinar populations. Expression of acinar markers (A) *Cpa1* or (B) *Try4* or ductal markers (C) *Krt18* or (D) *Krt19* overlaid on the UMAP of EYFP+ cells in Figure 2A. (E) Ridge plots of *Cpa1* and *Try4* expression in select acinar populations in Figure 2G, showing a loss of expression in ADM. (F) Mki67 expression either overlaid on the UMAP from Figure 2G or shown as a Ridge plot. (G) Ridge plot of *Pcna* expression. (H) S or G2M scores applied to the normal and injury-induced acinar populations featured in Figure 2G.

**Supplementary Figure 3.**
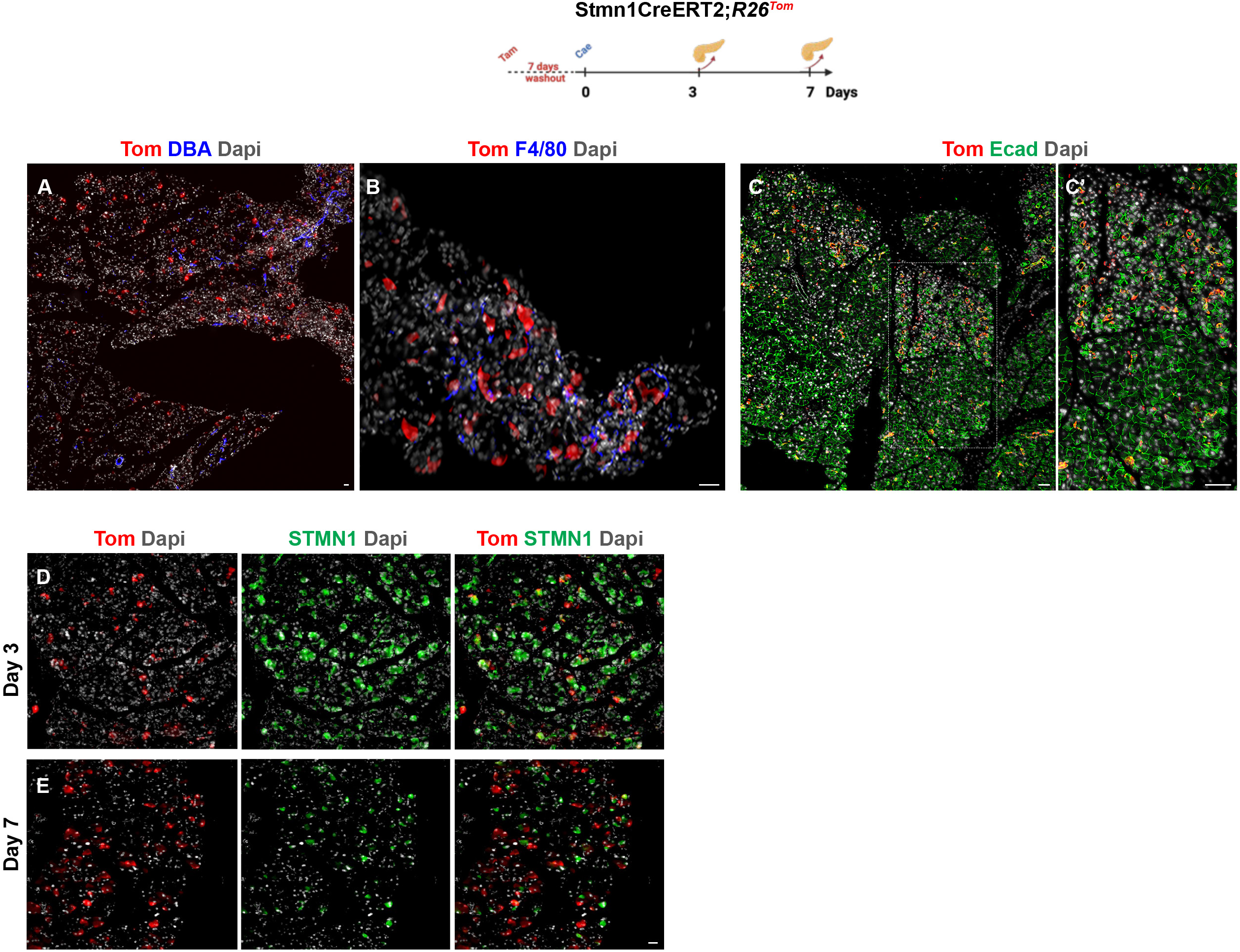
New onset of *Stmn*1 expression in acinar cells following injury. (A-C) Fluorescent imaging of Tom^+^ cells in SC*^Tom^* pancreas for detection of DBA (A), F4/80 (B) or E-cadherin (C, C’) showing focal expansion of Tom^+^ cells in acinar lobes that have been injured following caerulein treatment. (D, E) Fluorescent imaging of Tom^+^ cells in SC*^Tom^* pancreas for detection of STMN1 on days 3 (D) or 7 (E) following caerulein treatment. Scale bar 20μm.

**Supplementary Figure 4.**
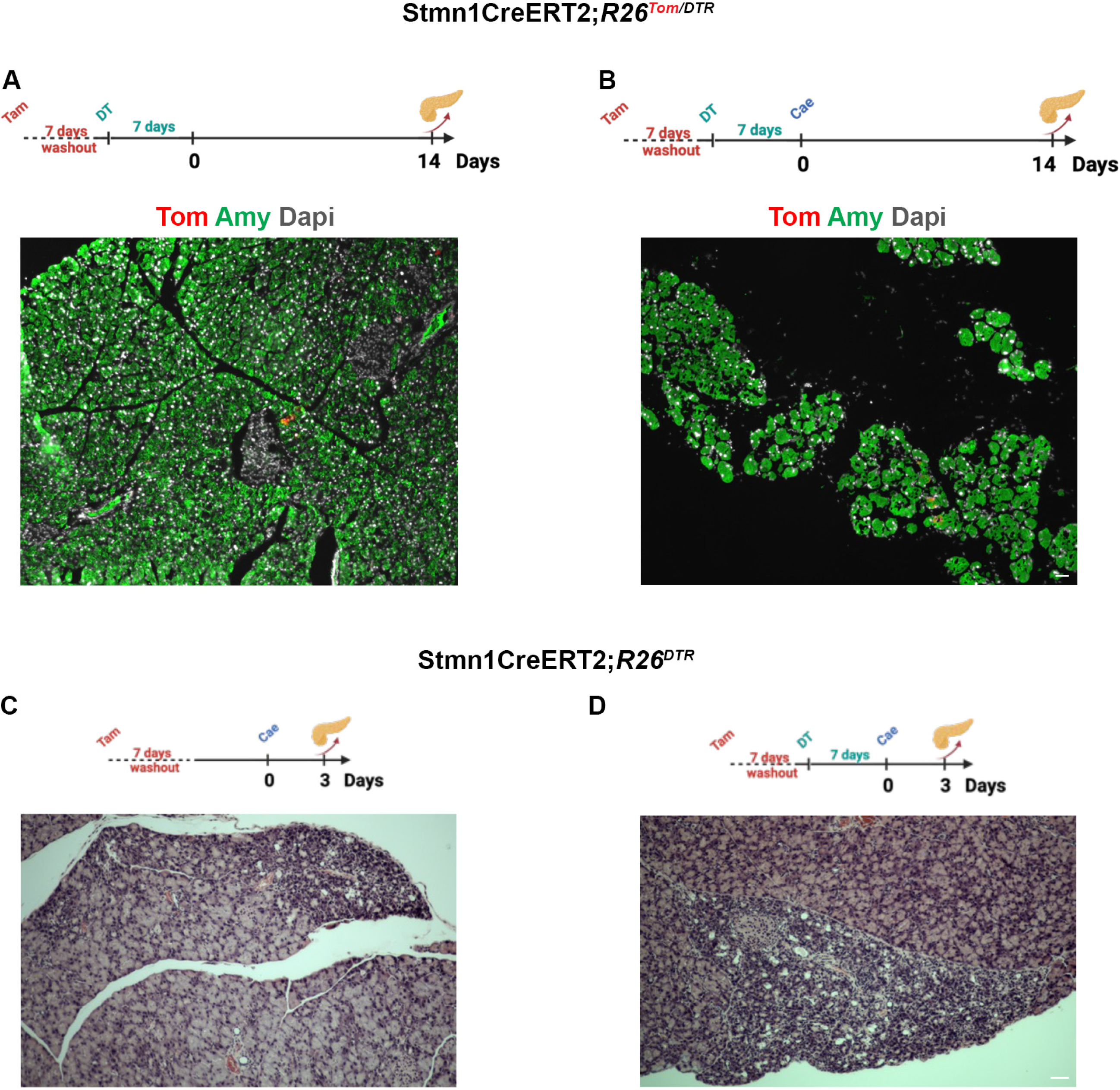
Stmn1-lineage is critical for acinar regeneration. (A, B) Representative images from pancreas sections of DT-treated SC*^Tom/DTR^* mice harvested 14 days after saline (A) or caerulein (B) treatments. (C, D) Representative data from pancreas sections of SC*^DTR^* mice harvested on day 3 post-caerulein without (C) or with (D) previous DT treatment. Scale bars 20μm.

**Supplementary Figure 5.**
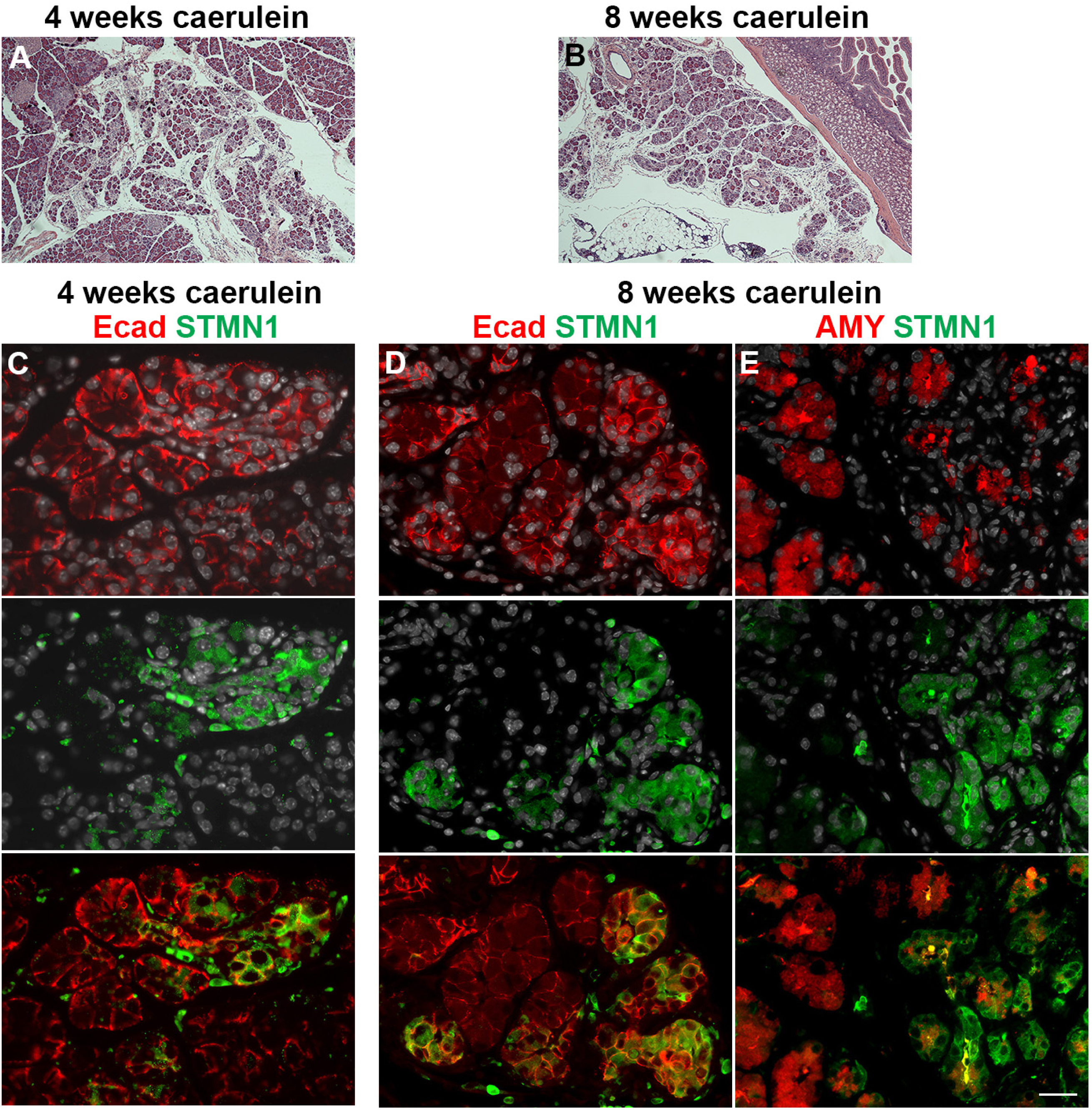
*Stmn1* expression during CP progression. (. A, B) H&E staining of tissues obtained from wild-type mice treated with caerulein for 4 (A) or 8 weeks (B) showing progression of CP. (C-E) Immuno-fluorescent staining for detection of E-cadherin and STMN1 (C, D) or amylase and STMN1 (E) 4-weeks (C) and 8-weeks (D, E) cohorts show expression of Stmn1 in ADMs and a subset of acinar cells. Note that in 8-weeks cohort (E), Stmn1 expression in acinar cells is associated with reduced or loss of amylase. Scale bar 20μm.

**Supplementary Figure 6.**
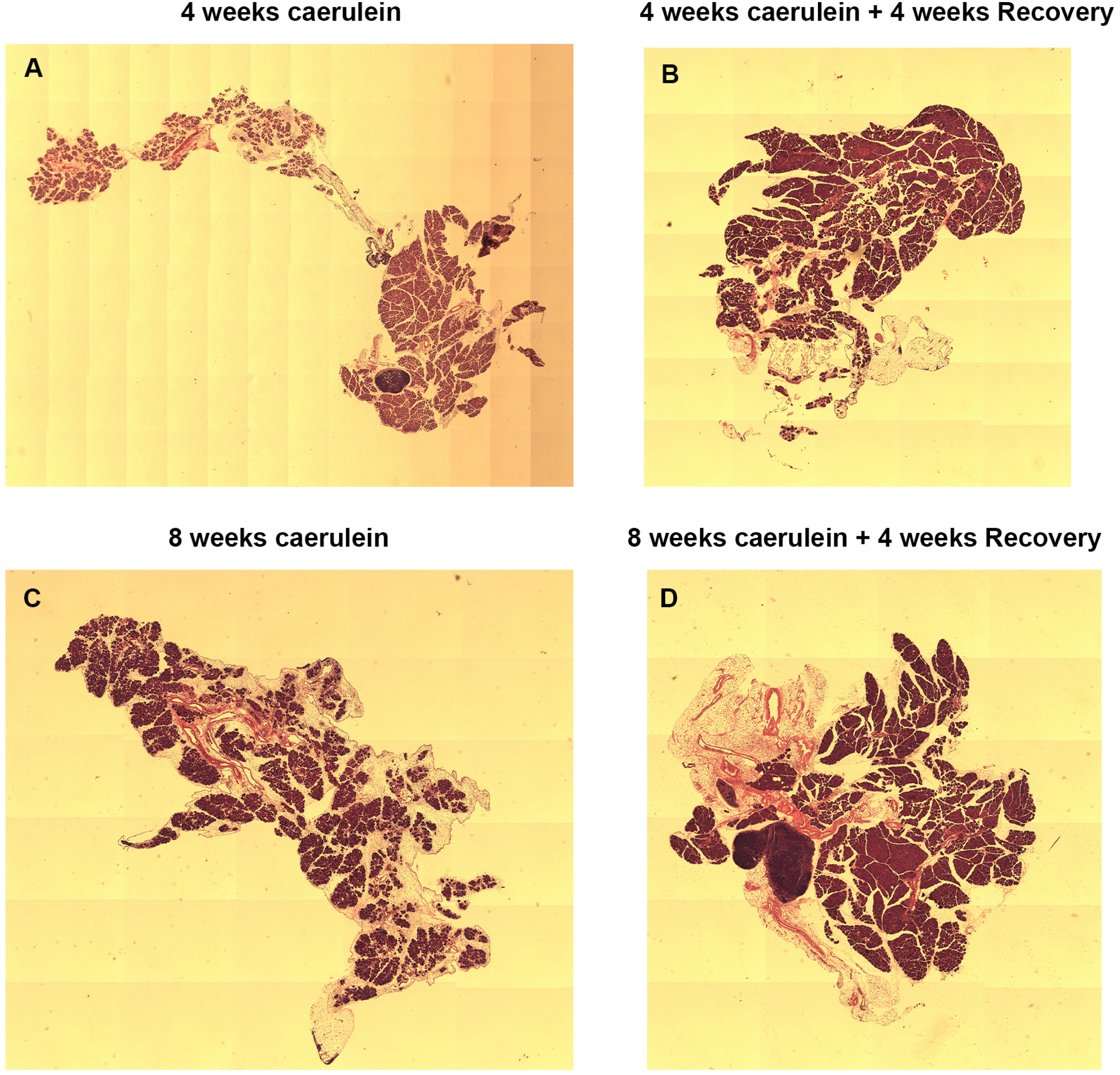
Acinar regeneration during CP progression. (A-D) Panoramic view of sections stained with hematoxylin and eosin from the pancreas of SC*^Tom^* mice harvested after 4 weeks caerulein treatment (A), 4 weeks caerulein treatment and 4 weeks recovery (B), 8 weeks caerulein treatment (C), or 8 weeks caerulein treatment and 4 weeks recovery (D).

**Supplementary Figure 7.**
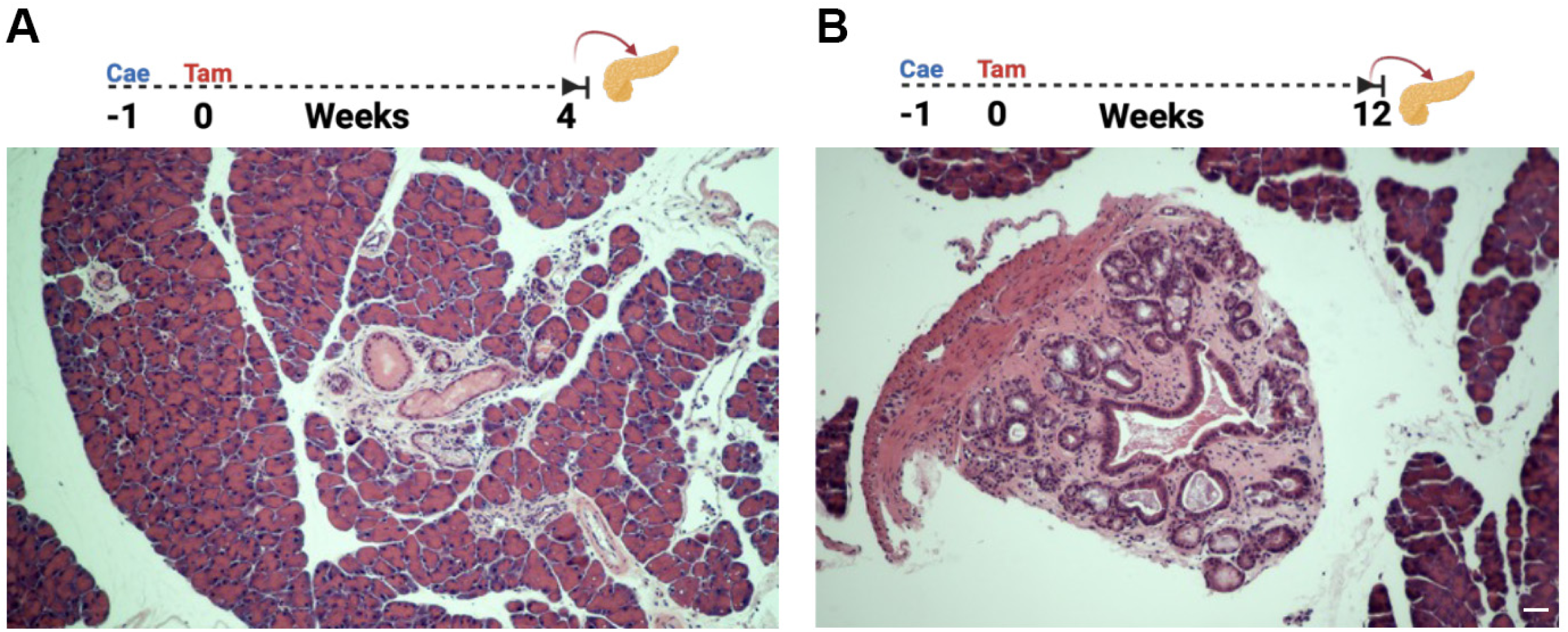
Cells with new onset of *Stmn1* expression can form PanINs. (A, B) Representative images from pancreas sections of SC*^Kras^* mice treated first with caerulein and harvested 4 (A), or 12 (B) weeks post-caerulein. Scale bar 20μm.

